# Widespread Autonomic Physiological Coupling Across the Brain-Body Axis

**DOI:** 10.1101/2023.01.19.524818

**Authors:** Taylor Bolt, Shiyu Wang, Jason S. Nomi, Roni Setton, Benjamin P. Gold, Blaise deB.Frederick, B.T. Thomas Yeo, J. Jean Chen, Dante Picchioni, R. Nathan Spreng, Shella D. Keilholz, Lucina Q. Uddin, Catie Chang

**Affiliations:** Department of Psychiatry and Biobehavioral Sciences, University of California Los Angeles, Los Angeles, CA, USA; Department of Biomedical Engineering, Vanderbilt University, Nashville, TN, USA; Department of Psychology, Harvard University, Boston, MA, USA; Departments of Electrical and Computer Engineering and Computer Science, Vanderbilt University, Nashville, TN, USA; Brain Imaging Center McLean Hospital, Harvard Medical School, Belmont, Massachusetts; Department of Electrical & Computer Engineering, Centre for Translational MR Research, Centre for Sleep & Cognition, N.1 Institute for Health and Institute for Digital Medicine, National University of Singapore, Singapore; Rotman Research Institute, Baycrest Health Sciences, Toronto, Canada; Department of Medical Biophysics, University of Toronto, Toronto, Canada; Institute of Biomedical Engineering, University of Toronto, Toronto, Canada; Advanced MRI Section, Laboratory of Functional and Molecular Imaging, National Institute of Neurological Disorders and Stroke, National Institutes of Health; Bethesda, MD, United States; Montreal Neurological Institute, Department of Neurology and Neurosurgery, McGill University, Montreal, QC, Canada; Emory University/Georgia Institute of Technology, Atlanta, GA, USA; Department of Psychology, University of California Los Angeles, Los Angeles, CA, USA

## Abstract

The brain is closely attuned to visceral signals from the body’s internal environment, as evidenced by the numerous associations between neural, hemodynamic, and peripheral physiological signals. We show that these brain-body co-fluctuations can be captured by a single spatiotemporal pattern. Across several independent samples, as well as single-echo and multi-echo fMRI data acquisition sequences, we identify widespread co-fluctuations in the low-frequency range (0.01 - 0.1 Hz) between resting-state global fMRI signals, neural activity, and a host of autonomic signals spanning cardiovascular, pulmonary, exocrine and smooth muscle systems. The same brain-body co-fluctuations observed at rest are elicited by arousal induced by cued deep breathing and intermittent sensory stimuli, as well as spontaneous phasic EEG events during sleep. Further, we show that the spatial structure of global fMRI signals is maintained under experimental suppression of end-tidal carbon dioxide (PETCO2) variations, suggesting that respiratory-driven fluctuations in arterial CO2 accompanying arousal cannot explain the origin of these signals in the brain. These findings establish the global fMRI signal as a significant component of the arousal response governed by the autonomic nervous system.

## Introduction

The communication of visceral signals from the body’s internal environment to the brain is essential for adaptive behavior. Central autonomic nuclei are closely connected to ascending arousal system nuclei that project diffusely to the cortex and subcortex, influencing wakefulness, affect and interoception (Satpute et al., 2019; Yackle et al., 2017). Pharmacological and electrical stimulation of central autonomic nuclei (e.g. solitary nucleus) produces widespread cortical EEG synchronization, suggesting a role for these structures in arousal and wakefulness fluctuations (Anaclet & Fuller, 2017; Dringenberg & Vanderwolf, 1998). Recent evidence suggests that widespread low-frequency (0.01-0.1 Hz) signals recorded from functional magnetic resonance imaging (fMRI), referred to as the ‘global signal’ (Liu et al., 2017), may also be a component of the central and peripheral nervous system arousal response (Liu et al., 2018; Özbay et al., 2019; Raut et al., 2021).

The ‘global signal’ is the most prominent spatiotemporal pattern observed in resting-state fMRI signals and accounts for a wide variety of previously observed features of these signals, including functional connectivity gradients (Bolt et al., 2022; Raut et al., 2021). Global fMRI signals are prominent during changes of arousal, as measured by electrophysiology, including EEG (Özbay et al., 2019) and local field potentials (Liu et al., 2018), as well as changes in peripheral physiology, such as pupil dilation and peripheral skin vascular tone (Gu et al., 2022; Özbay et al., 2019; Raut et al., 2021). Özbay et al. (2019) have shown that fluctuations in arousal during sleep co-occur with simultaneous fluctuations in global fMRI signals and peripheral skin vascular tone, an indicator of sympathetic-mediated vasoconstriction.

This study seeks to demonstrate the degree of coordination between global fMRI, EEG and autonomic signaling across multiple ANS effector organs. Using multiple independent samples of multi-modal fMRI, EEG and peripheral physiological recordings acquired during resting state, we demonstrate that a single, low-dimensional projection captures a major axis of covariation between global brain fluctuations and widespread peripheral physiological dynamics, including pulmonary (respiratory variability), cardiovascular (heart rate variability), exocrine (skin conductance) and smooth muscle (peripheral vascular tone and pupil diameter) systems.

Spontaneous variations in autonomic arousal (K-complexes) during sleep, as well as direct manipulation of autonomic activity via deep breathing and sensory stimulation, induced a similar pattern of co-fluctuations between the brain and body. Experimental suppression of spontaneous variations in end-tidal carbon dioxide (PETCO2) accompanying arousal demonstrated that arterial CO2 could not explain the origin of these global brain fluctuations.

Taken together, these findings provide novel evidence for the role of sympathetic-mediated vasoconstriction and ascending arousal system projections in governing spontaneous global fluctuations commonly observed in fMRI. This provides insight into long-standing debates regarding the origins and functional significance of global fMRI signals (Liu et al., 2017; Uddin, 2020).

## Results

### Global fMRI Fluctuations Are Associated with Systemic Physiological Changes

Global fMRI fluctuations were extracted via the first principal component (PC1) of whole-brain fMRI time courses (Bolt et al., 2022). The spatial and spatiotemporal distribution of the first PC derived from PCA and CPCA, respectively, is displayed in **Figure 1A**. Examination of the spatiotemporal structure of global fMRI studies are analyzed further below (**Figure 3**).

**Figure 1.**
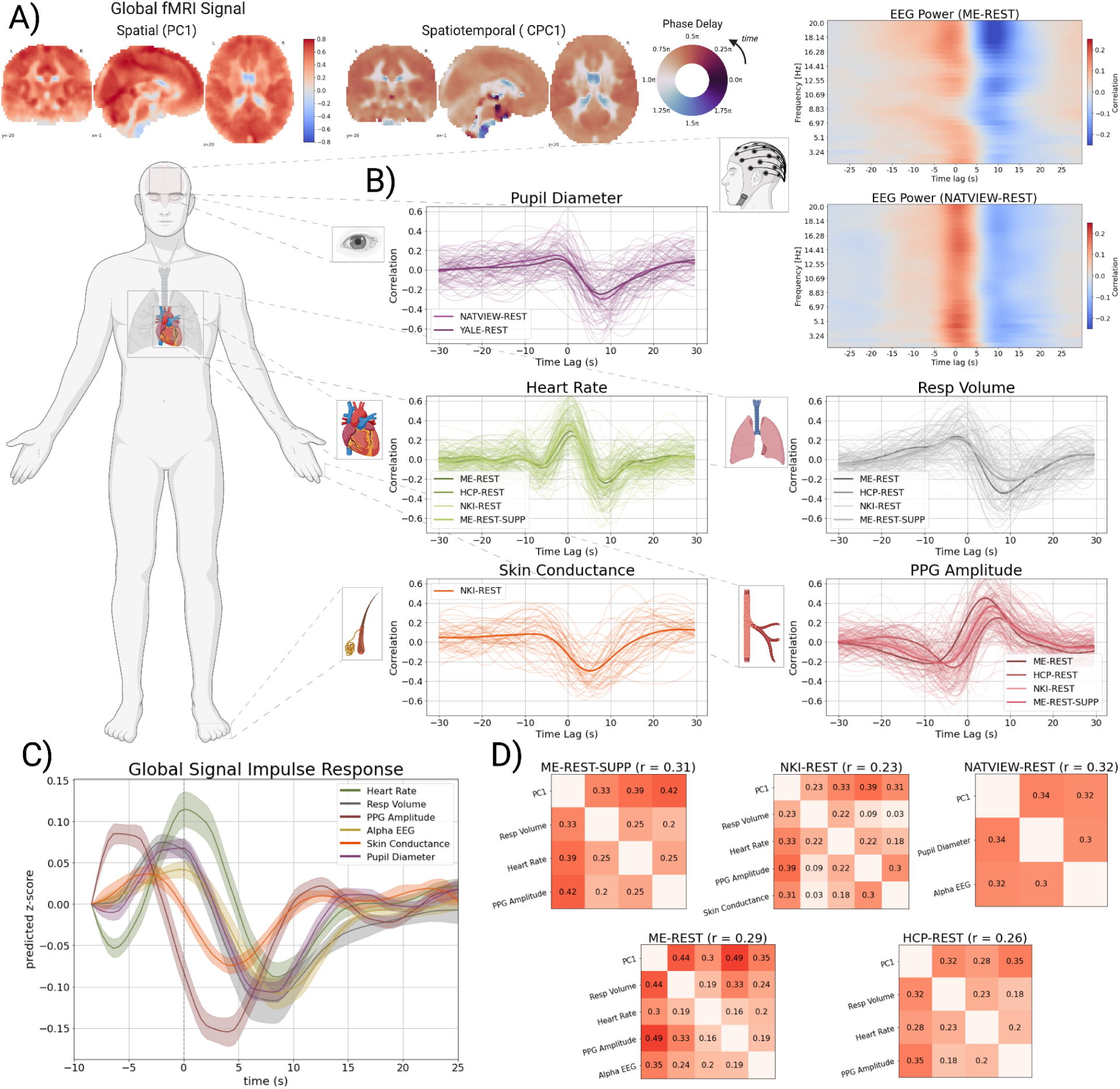
Global fMRI Fluctuations are Associated with Systemic Physiological Changes. The cross-correlation between the time courses of the global fMRI signal (first principal component; PC1) and multiple physiological signals. **A)** The spatial weights of the first principal component (PC1; top), and the phase delay map of the first complex principal component (CPC1; bottom) from the ME-REST dataset. The phase delay map of the first complex principal component encodes the time-delay (in radians) between voxels within the component. Because phase delay is measured in radians (0 to 2pi) they are displayed with a circular color map. **B)** Cross-correlation plots of each physiological signal with the global fMRI time course (PC1). The cross-correlation is defined as the correlation coefficient between *x*_*t*+*i*_ and *y*_*t*_ where *t* is the time index, *i* is ± 30 secs (i.e. the index along the x-axis of the plots), and *x* is the global fMRI signal and *y* is the physiological signal. Strong correlation at a positive time lag (i.e. positive *i* index) indicates that the global fMRI signal lags or follows the physiological signal, while strong correlation at a negative time lag (i.e. negative *i* index) indicates that the global fMRI signal leads the physiological signal. Within a dataset, all subject-level cross-correlations are displayed in lighter, more transparent color, while the mean signal across subjects is displayed in a darker color. Each dataset is displayed in a separate shade of the same color. Cross-correlations between global fMRI signals and wavelet filtered EEG power signals (Morlet wavelet; number of cycles = 15) for the ME-REST and NATVIEW-REST datasets are displayed as a heat map in the top panel. **C)** Estimated impulse response functions of the global fMRI signal in response to an impulse of each physiological signal. The global signal response to PPG amplitude is inverted for consistency with the other physiological signals that exhibit a positive peak at positive lags of the global fMRI signal (see **Panel B**). Negative time points are included to visualize the initial positive peak of the global fMRI signal response. Standard errors are calculated from cluster bootstraps (N_bootstraps_ = 500) **D)** The pairwise correlations between all physiological signals (including the global fMRI signal; PC1) in the first canonical component of the MCCA analysis. The average pairwise correlation is displayed beside the title of each correlation matrix. The pairwise correlations of the first canonical component indicate a strong, joint co-fluctuation between all physiological signals and the global fMRI signal across all datasets.

To examine the association of global fMRI signals with peripheral physiological signals, we computed a single global fMRI signal from the first PC time course, such that positive signal fluctuations reflected positive fMRI signal in the cortex and negative signal in CSF compartments (**Figure 1A**). Analysis of six separate resting-state fMRI datasets, several collected in the laboratories of co-authors (ME-REST, ME-REST-SUPP) and others conducted using publicly available datasets (HCP-REST, NATVIEW-REST, NKI-REST, and YALE-REST), shows that global fMRI fluctuations covary with a multitude of peripheral physiological signals (**Figure 1B**), including respiratory volume, heart rate, PPG amplitude (a measure of peripheral vascular tone), skin conductance, and pupil diameter. Analysis of the relationship between surface EEG power fluctuations and global fMRI signals in two simultaneous EEG-fMRI datasets (ME-REST, NATVIEW-REST) revealed prominent co-fluctuations between global fMRI signals and neuronal oscillations across a wide band of frequencies (2 - 20Hz). Most cross-correlation plots exhibit two distinct peaks for all signals, indicating temporal associations between physiological signals and the global fMRI signal at two distinct time lags: 1) a negative or zero time lag (0 to ∼ 5 secs) of the global fMRI signal, indicating that the physiological signal follows the global fMRI signal, and 2) a positive lag (0 to ∼10s) of the global fMRI signal, indicating that the physiological signal precedes the global fMRI signal.

To examine the temporal dynamics of the global fMRI signal associated with these physiological signals, we estimated impulse response functions (see *Methods and Materials*; Chang et al., 2009) of the global fMRI signal in response to an impulse of each physiological signal: ME-REST (PPG amplitude, Alpha EEG power, and respiratory volume), NATVIEW-REST (pupil), and NKI-REST datasets (skin conductance) (**Figure 1C**). The dynamics of the global fMRI signal exhibit a consistent pattern across all physiological signals - a positive peak followed by a stronger and more prolonged negative peak before returning to baseline.

The cross-correlation results establish pairwise relationships between global fMRI signals and physiological signals, but do not provide definitive evidence of joint co-fluctuations between all pairs of signals across time. To establish the latter, we performed a cross-decomposition between all pairs of signals and their lags via multi-set canonical correlation analysis (MCCA; see *Methods and Materials*). Our aim was to examine whether the joint co-fluctuations between all signals (and their time-lags) can be extracted in a single latent dimension. Note, though the relationship between global fMRI and EEG power is broadband (see **panel B**), we included alpha (8-12Hz) EEG power as a single signal from the EEG due to previous reports of a relationship between alpha power and global fMRI signals (Yuan et al., 2013).

Application of MCCA to five resting-state fMRI datasets demonstrates that the first canonical component (**Figure 1D**), representing the latent component with the maximal average pairwise correlation between all pairs of signals, captures moderate pairwise correlations between all signals, including peripheral physiology, neuronal (alpha EEG power) and global fMRI (PC1) signals (e.g ME-REST: ***r̄*** 0.31, p = 0.001). Further, the global fMRI signal exhibits the strongest pairwise correlations among all signals within the first canonical component across datasets. Time lags between signals in the first canonical component are presented in **Supplementary Figure 10**.

### Autonomic Arousal Elicits Brain-Body Co-Fluctuations

If the ANS is the source of these co-fluctuations across brain and body, arousal induction should produce a similar pattern of co-fluctuations as that observed at rest (**Figure 1**). To provide empirical support for this hypothesis, we performed event-related averaging on fMRI, EEG and physiological signals from 1) a cued deep inhalation task (ME-TASK), 2) a cued reaction time task (ME-TASK-CUE), and 3) K-complex onsets for participants who fell asleep during the resting-state session of the ME-REST dataset. The cued deep inhalation task (ME-TASK) was a sparse event-related design with individual cues separated by one to two minutes, allowing for an estimation of the physiological impulse response to isolated inhalations without overlapping responses. The cued reaction time task was a sparse event-related design similar to the deep inhalation task, but a button response was performed in response to the cue. Consistent with a previous study (Özbay et al., 2019), we also chose to examine physiological responses in and around K-complex onsets during stage II sleep, which was found in a subset of participants of the ME-REST dataset. The K-complex is a characteristic high-amplitude EEG event that occurs predominantly in stage II sleep, and reflects phasic arousal events from internal (interoceptive) or external stimuli (Cash et al., 2009; Halász et al., 2004). K-complex annotations were performed in a semi-automated fashion on the EEG time courses of the ME-REST dataset (see *Methods and Materials*) and manually reviewed for accuracy.

Global fMRI signals (PC1) exhibit large amplitude fluctuations to isolated deep inhalations (ME-TASK) (**Figure 2A**). Consistent with the shape of the respiration response function described by Birn et al. (2008), global fMRI signals exhibit a bimodal response to inhalation, with an early positive increase (∼4s) followed by a prolonged undershoot reaching its trough around ∼14s. In response to isolated deep inhalations, physiological signals exhibit temporal dynamics consistent with increased ANS activity (**Figure 2A**), and lead-lag timing that is similar to that observed at rest (**Figure 1**). An increase in heart rate, broadband EEG power (**Figure 2B**), and peripheral vasoconstriction (decrease in PPG amplitude) is observed around the same time as the early positive peak (∼4s) of the global fMRI signal.

**Figure 2.**
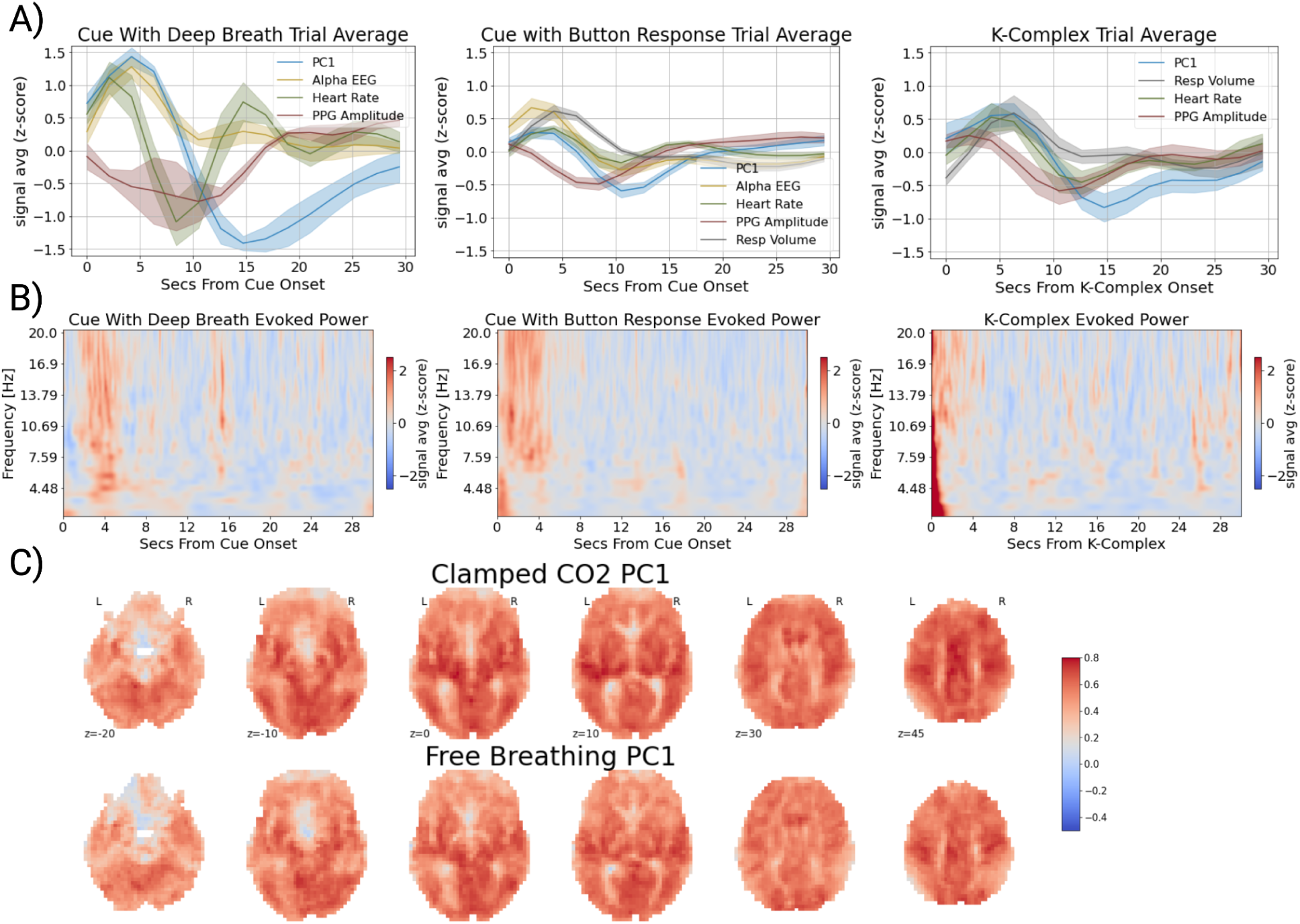
Arousal Changes Generate Joint fMRI-Physiological Co-Fluctuations. A) Event-related averages of physiological signals and global fMRI signals in response to inhalation with an auditory cue (left; N_trials_ = 76; N_subj_ = 6), an auditory cue with button response (middle; N_trials_ = 254; N_subj_ = 12) and K-complexes during sleep (right; N_trials_ = 64; N_subj_ = 7). Averages are displayed in z-score units. Standard error bars for the physiological time courses are constructed via cluster bootstrapping (i.e. resampling at the subject level; n_bootstraps_ = 500). The strongest amplitude responses were observed in inhalation trials. A similar pattern of responses were observed in response to auditory cues with button responses and around spontaneous K-complexes during sleep, but with smaller magnitude. **B)** Evoked EEG power averages to inhalation with an auditory cue (left), an auditory cue with button response (middle) and K-complexes during sleep (right) were constructed through trial-averaging of Morlet wavelet filtered power signals within subjects (see Methods), followed by group-averaging across subject-trial averages. **C**) The spatial weights of the first principal component (PC1; global fMRI signal) estimated from a free breathing and ‘clamped’ CO2 condition.

The physiological response to a paced inhalation/breath hold task was also examined (NKI-TASK). As opposed to the sparse-event design of the inhalation task, this task was organized in sequential blocks with deep inhalations/exhalations followed by a prolonged breath hold (18s). This task structure was found to also elicit strong amplitude fluctuations in peripheral physiology and global fMRI signals with peak/trough timings largely consistent with the deep inhalation task (**Supplementary Figure 4**).

To understand if this spatiotemporal pattern of physiological dynamics is not specific to breathing, but to arousal induction more broadly, we examined physiological dynamics during a simple reaction time task, where button presses were performed in response to intermittent auditory cues. While smaller in magnitude, a similar spatiotemporal pattern was observed in button responses (**Figure 2A**), though the undershoot of the global fMRI signal occurred several seconds earlier in the button response compared with the deep inhalations.

We also examined whether spontaneous autonomic arousals produced similar physiological responses as that observed in resting-state, external stimuli, and deep breathing. Event-related averages of physiological and global fMRI signals around K-complex onsets revealed the same pattern of physiological response as that observed in deep breathing, as well as that observed spontaneously during rest (**Figure 2A-B**).

### Global fMRI Signals Under Clamped CO2 Conditions

One potential source of global fMRI fluctuations in response to arousal induction is changes in cerebral blood flow due to levels of arterial CO2, a vasodilator (Battisti-Charbonney et al., 2011; Wise et al., 2004). As has been shown in the current study and others (e.g., Gu et al., 2022), changes in respiratory volume (respiratory rate and depth) can induce, as well as accompany, arousal state changes. Sustained changes in respiratory volume modulate the level of arterial CO2, thereby potentially increasing cerebral blood flow (Birn et al., 2006; Wise et al., 2004). We sought to test if spontaneous fluctuations in arterial CO2 served as the sole driver of the global fMRI pattern observed in this study. To do so, we examined the existence and spatial structure of global fMRI signals under both free breathing and under a ‘clamped’ CO2 condition, where end-tidal CO2 (PETCO2) was ‘clamped’ to the average level of each participant (Golestani & Chen, 2020).

The spatial structure of the global fMRI signal was estimated with the spatial weights of the first principal component and compared across free breathing and ‘clamped’ CO2 conditions (**Figure 2C**). The spatial structure of global fMRI signals was found to be nearly identical between free breathing and clamped CO2 conditions (correlation r = 0.95 of the spatial weights between the two conditions). Further, global fMRI signals were similarly present in both conditions, as reflected in the explained variance estimates of the first principal component between the two conditions (PC1_free_ r^2^ = 0.28; PC1_clamped_ r^2^ = 0.25; **Supplementary Figure 11**).

Consistent with a previous report comparing overall time-lag structure of global fMRI signals between clamped and free breathing conditions (Kish et al., 2023), we found that the spatiotemporal structure of global fMRI signals was similarity maintained in clamped CO2 conditions (**Supplementary Figure 5**).

### Spatiotemporal Dynamics of the Global fMRI Signal

To examine the spatiotemporal dynamics of the global fMRI signal and its anatomical distribution, we visualized brain maps at selected time points of a temporal reconstruction of the global fMRI signal through CPCA (see *Methods and Materials*; **Figure 3C**). An alternative approach of extracting the spatiotemporal pattern from voxel-wise cross-correlations with PPG amplitude, the physiological signal with the strongest association with the global fMRI signal, yielded the same pattern (**Figure 3B**). Multi-echo fMRI data (ME-REST-SUPP) was used to isolate blood-oxygen level dependent (BOLD) signal changes in the global fMRI signal via multi-echo independent component analysis (ME-ICA) denoising.

**Figure 3.**
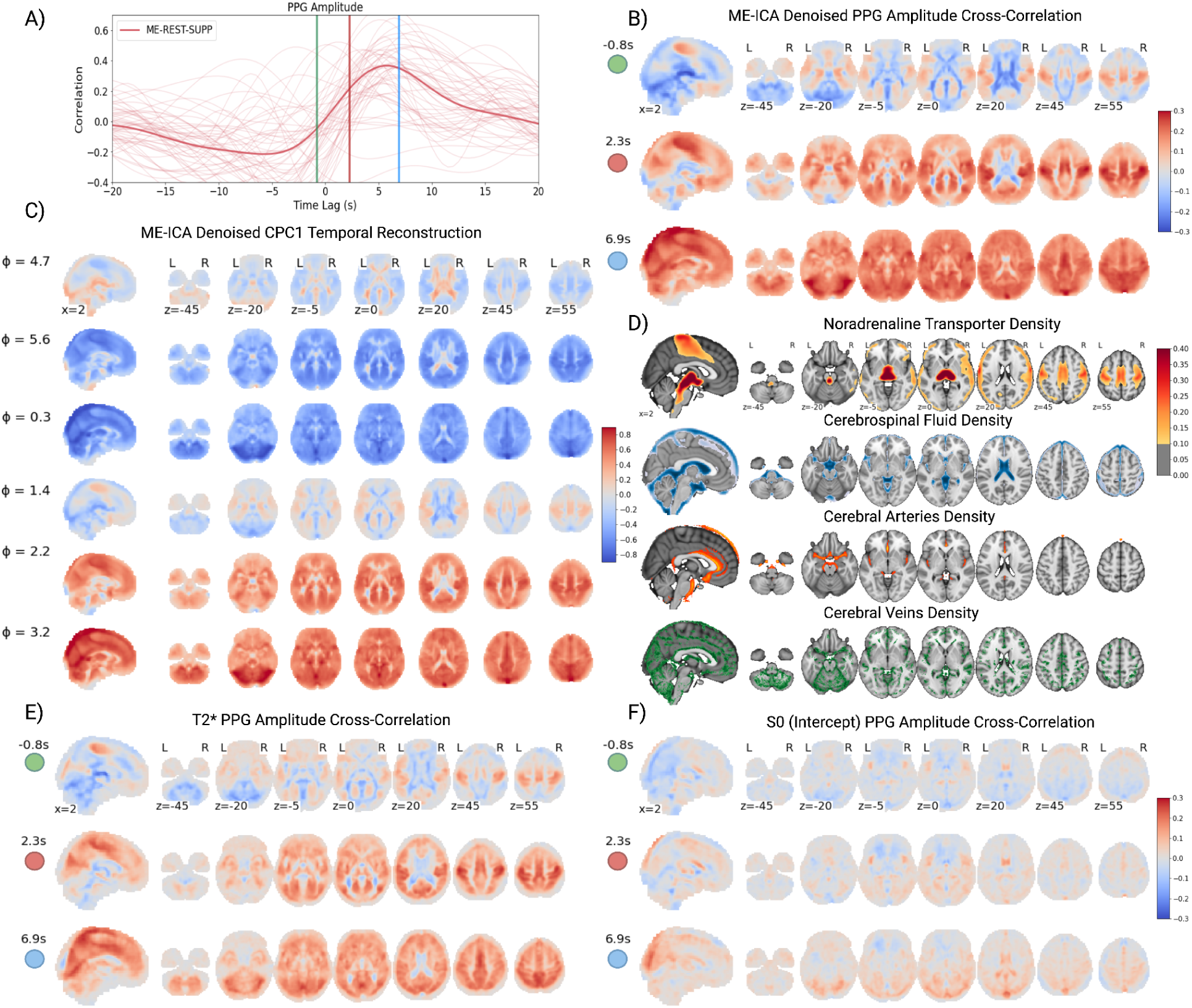
Spatiotemporal Dynamics of the Global fMRI Signal. A) Subject- (thin lines) and group-averaged (thick lines) cross-correlation plot of the global fMRI signal with PPG peak amplitude for the ME-REST-SUPP dataset. **B)** Due to the strong association between PPG peak amplitude and global fMRI signals, voxel-wise cross-correlations between PPG peak amplitude and fMRI signals were presented at select time points (green, red, blue) of the cross-correlation curve to illustrate the temporal dynamics of the global fMRI signal. **C)** The spatiotemporal reconstruction of the global fMRI signal from CPCA on the ME-REST-SUPP optimally combined (ME-ICA denoised) time courses displayed at select phases (in radians - 0 to 2pi). Note, the phase (ɸ) is arbitrary and ‘wraps around’ in the interval [0, 2pi], such that ɸ=0 or ɸ=2pi does not mark the beginning or end of the spatiotemporal pattern, respectively. **D)** First (from top): Noradrenergic transporter (NAT) density (Ding et al., 2010), second: thresholded (> 0.6) cerebrospinal fluid probability map (Lorio et al., 2016), third: thresholded (>0.6) cerebral veins probability map (Huck et al., 2019) and fourth: thresholded (>0.1) cerebral arteries probability map (Mouches & Forkert, 2019). **E)** The cross-correlation of multi-echo estimated **E)** T2* (slope) and **F)** S0 (intercept) time courses with PPG amplitude at the selected time points in **panel A**.

CPCA models the global fMRI spatiotemporal pattern as cyclical with mirrored globally positive (ɸ=0.3) and negative (ɸ=3.2) peaks (**Figure 3C**). For clarity, we define the onset of the spatiotemporal pattern at ɸ=4.7, corresponding to the pattern of fMRI signals observed at PPG amplitude drops (t=0, **Figure 3A**; **Figure 3B** - Top Panel). The global fMRI signal exhibits a stereotypical pattern: 1) an early decrease in fMRI signals in the sensorimotor and visual cortices concomitant with an increase of signals overlapping the white matter, CSF compartments and large cerebral arteries (ɸ=4.7, **Figure 3C**; t=-0.8s, **Figure 3B**) 2) followed by a decrease in fMRI signals across the gray and white matter (ɸ=5.6, **Figure 3C**; t=2.3s, **Figure 3B**) and 3) finally a delayed decrease in fMRI signals in the large cerebral veins (ɸ=0.3, **Figure 3C**; t=6.9s, **Figure 3B**).

The presence of the global fMRI signal during changes in arousal (**Figure 2**) and its association with pupil diameter (**Figure 1**) suggests increased signaling of cortical noradrenergic projections from ascending arousal stimuli during global fMRI fluctuations. To examine whether any components of the global fMRI signal are potentially related to noradrenergic signaling from ascending arousal nuclei, we compared the spatiotemporal pattern to an atlas of noradrenaline transport (NAT) density (binding potential) derived from positron emission tomography (Ding et al., 2010; Markello et al., 2022) (**Figure 3D).** In the cerebral cortex, NAT density is most prominent in the sensorimotor area, overlapping the same region with early fluctuations of the global fMRI signal.

To explore the biophysical underpinnings of the global fMRI signal, we characterized signal decay properties at select time points of the cross-correlation of fMRI signals with PPG amplitude. Specifically, we estimated echo-time (TE) dependent and TE-independent signals (see *Methods and Materials*) by ordinary least squares fit of a monoexponential decay curve to the consecutively acquired echos for each volume, 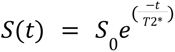, where T2* (slope from a log-linear fit) and S0 (intercept) estimate the TE-dependent and TE-independent signal, respectively. T2* estimates on a volume-wise basis reflect noisier estimates of BOLD signal changes relative to the ME-ICA denoised signals (**Figure 3C**), but yield a similar global fMRI spatiotemporal pattern (**Figure 3E**). S0 and T2* maps were found to exhibit opposing changes in CSF compartments (including the ventricles and areas along the midline and insula) and posterior draining veins (**Figure 3F**). The opposing changes in T2* and S0 signals in these regions are suggestive of local blood volume changes with little to no blood oxygenation (BOLD) changes, in line with a bicompartmental model of fMRI signal composed of CSF and blood (Bianciardi et al., 2011).

## Discussion

The composition and functional significance of the global fMRI signal is widely debated (Liu et al., 2017; Uddin, 2020). This study characterizes a large-scale pattern of co-fluctuations between global fMRI brain signals and sympathetic-mediated physiological fluctuations in humans. This pattern of co-fluctuations is replicated across multiple independent samples of multi-modal fMRI, EEG and peripheral physiology recordings. It is widespread across the body and entire nervous system, involving the brain, heart, lungs, exocrine and smooth muscle systems. It is also linked to changes in arousal state induced via deep breathing and sensory stimulation, as well as spontaneous fluctuations in arousal observed during sleep.

The co-fluctuations between global brain signals and peripheral physiology exhibit a stereotypical sequence. The initial positive increase in the global fMRI spatiotemporal pattern is accompanied by roughly simultaneous increases in heart rate, peripheral vascular tone, respiratory volume and broadband EEG power, followed by a drop in the amplitude of the PPG signal. The increase and subsequent decrease in heart rate around simultaneous peaks of respiratory volume is consistent with gating of parasympathetic outflow (via the vagus nerve) during inspiration and its relaxation during exhalation, known as respiratory sinus arrhythmia (Yasuma & Hayano, 2004). The drop in PPG amplitude is indicative of vasoconstriction of peripheral blood vessels by sympathetic outflow (Khoo & Chalacheva, 2019). While pupil diameter and tonic skin conductance signals were only recorded in resting-state conditions, impulse response and cross-correlation analyses (**Figure 1 B,C**) provide potential insights into their dynamics during evoked conditions; these suggest an early increase in pupil diameter followed by an increase in skin conductance. An increase in pupil diameter is consistent with inhibition of parasympathetic-mediated outflow to the iris constrictor muscles and excitation of sympathetic-mediated outflow to the iris dilator muscle (Marumo & Nakano, 2021), and changes in skin conductance are consistent with sympathetic-mediated changes in sudomotor activity.

We observed that experimentally-induced or spontaneous variance in ANS activity reproduces the same sequence of physiological and global brain activity observed at rest. Here, experimental manipulation of ANS activity was conducted through cued deep breathing and sensory stimulation, but other autonomic challenges may produce a similar effect. Respiratory variation is a strong modulator of peripheral ANS activity and central arousal. Modulation of sympathetic outflow in the periphery through changes in respiratory rate and depth is a well-established finding in human physiology studies (Häbler et al., 1994). Further, neuronal populations within the medulla responsible for respiratory rhythm generation (preBötzinger complex) directly project to, and modulate, activity of noradrenergic neurons in the locus coeruleus. While smaller in magnitude, intermittent auditory-cued button responses reproduced the same sequence of physiological and global brain activity observed during deep breathing. The same spatiotemporal sequence was also observed around K-complex onsets during sleep, consistent with a previous report (Özbay et al., 2019) that found increased global fMRI-physiological coupling around K-complex onsets during the NREM2 sleep stage. The coupling of ANS-related physiological signals and K-complex onsets are generally understood to reflect the central activation of sympathetic nervous system outflow (de Zambotti et al., 2018)

Spatiotemporal modeling of the global fMRI signal via CPCA (**Figure 3**; **Supplementary Figure 1; Supplementary Movies 1, 2**) revealed a characteristic time-lag pattern, where early signal changes in sensorimotor and visual cortices propagate across the gray matter with concomitant opposing signal changes in white matter, CSF compartments and cerebral arteries. Cross-correlations of whole-brain fMRI signals with PPG amplitude, the physiological signal with the strongest association with the global fMRI signal (**Figure 1B; 1D**), yielded the same spatiotemporal pattern as the CPCA approach. The sign and timing of the cross-correlation of PPG amplitude and whole-brain fMRI signals are in agreement with previous studies (Özbay et al., 2018).

Several features of the global fMRI spatiotemporal pattern are unlikely related to local neurovascular coupling mechanisms. First, simultaneous with drops in PPG amplitude, negative signals are observed in the gray matter and positive signals in the white matter. Several seconds (∼5s) following drops in PPG amplitude, negative fMRI signals are observed in the white matter. The early increase followed shortly by a decrease in white matter fMRI signals is likely due to the mechanism proposed by Özbay et al. (2018): differences in arterio-venous transit time between the pial vessels innervating gray matter and deep medullary vessels innervating the white matter. Decreased cerebral arterial blood flow results in temporally overlapping decreases of blood volume and oxygenation in pial veins of the gray matter, producing a net decrease in the fMRI signal due to the shorter arterio-venous transit time of oxygenated blood. In the deep medullary veins, however, a lag in arterio-venous transit time results in a temporary mismatch between blood volume and oxygenation fluctuations, such that early blood volume decreases precede oxygenation decreases resulting in a net increase in fMRI signal during this lag, eventually followed by a net decrease in fMRI signal.

Second, the negative gray-matter fMRI signal changes accompanying PPG amplitude drops also occurred together with positive signals in CSF compartments and large cerebral arteries. Early opposing signal changes in CSF compartments relative to the gray matter have been previously observed in studies of cerebrovascular reactivity (Bright et al., 2014; Thomas et al., 2013). Periventricular white matter and ventricle regions are susceptible to partial volume artifacts from nearby blood vessels. During vasodilation of nearby blood vessels, the increase in blood volume reduces CSF volume, inducing a net decrease in the fMRI signal (Thomas et al., 2013). This is due to the brighter CSF signal (long T2) being displaced by darker blood signal (shorter T2). During PPG amplitude drops, positive signals are observed in the CSF compartments suggesting the opposite response - a decrease in blood volume with increased CSF volume. Similar partial volume effects may explain opposing signals observed in large cerebral arteries relative to gray matter during PPG amplitude drops. Cerebral arteries with positive fMRI signals during PPG amplitude drops (e.g. basilar artery, internal carotid and posterior cerebral artery) overlap with areas of high CSF density (**Figure 3D**), suggesting decreases in arterial blood volume (and increases of CSF volume) may result in positive fMRI signal in these areas. Alternatively, opposing signals in the cerebral arteries may arise from partial-volume effects between arterial blood and surrounding tissue, such that blood volume fluctuations change the relative size of the intravascular component contribution to the signal (versus the extravascular component), resulting in changes in phase homogeneity within a voxel (Zhong et al., 2024).

The hypothesis that blood volume fluctuations are the source of positive signals in the CSF compartments and cerebral arteries is further supported by our T2* and S0 (initial signal intensity) modeling with multi-echo fMRI data (**Figure 3F**). Simultaneous with drops of PPG amplitude, positive T2* and negative S0 signal fluctuations are observed in areas of high CSF density. Modeling results (Bianciardi et al., 2011) have shown that these opposing signal changes in T2* and S0 signals are consistent with decreases in blood volume. It is important to note that our modeling of signal decay curves was limited to three echos and curve fits were likely to be noisy at the subject-level, and should be further confirmed in data collected with more echos.

Another possible source of the opposing signal changes in the CSF compartments relative to gray matter is the presence of CSF inflow effects from glymphatic pulsation mechanisms (Bohr et al., 2022) that have been previously observed in fMRI signals during states of drowsiness/sleep (Fultz et al., 2019; Gonzalez-Castillo et al., 2022) and autonomic arousal (Picchioni et al., 2022; Wang et al., 2022). Further, these studies have observed that CSF inflow signals are anti-correlated with gray matter fluctuations, an observation also found in our analyses. However, these spatiotemporal dynamics were observed in multi-echo denoised signals (**Figure 3B**) and directly estimated T2* signals (**Figure 3E**), where inflow effects have been (presumably) attenuated. In fact, in some cases, anti-correlated signals in the CSF compartments and white matter in the global fMRI signal are more prominent in multi-echo denoised signals (**Supplementary Figure 2**), suggestive of BOLD contrast changes in these areas. While BOLD signal changes in ventricles may be unexpected, the choroid plexus, a specialized epithelial tissue that lines the ventricles, is heavily vascularized and may be the source of hemodynamic effects in these areas. However, the presence of residual inflow effects cannot be conclusively ruled out.

Third, following the early decrease of signal in sensorimotor and visual cortices simultaneous with drops in PPG amplitude, large amplitude decreases are observed in and around cerebral veins (e.g., inferior and superior sagittal sinus, straight sinus, and transverse sinus). These are suggestive of the propagation of blood flow or volume changes to large draining veins from the parenchyma (brain tissue). This propagation is visible in both the phase delay maps from CPCA (**Figure 1A**) and movies of the spatiotemporal progression of global fMRI signal (**Supplementary Movie 1; 2**). The prominence of late vein effects in the global fMRI signal does not uniquely determine its upstream mechanism, as such effects would be present in upstream systemic vascular (e.g. vasoconstriction) or local neurovascular mechanisms (Turner, 2002). In fact, a delayed onset of fMRI signals in and around large cerebral veins distal to the site of neural activation is regularly observed in task-fMRI experiments (Kay et al., 2020; Sheth et al., 2005).

The spatiotemporal changes in fMRI signals just described are consistent with several ‘upstream’ causal mechanisms. An often-cited ‘upstream’ causal mechanism for global fMRI signals is respiratory-driven changes in arterial CO2 concentration (Wise et al., 2004). Changes in breathing depth and rate and respiratory reflexes occur frequently in resting-state fMRI scanning sessions. Power and colleagues (Lynch et al., 2020; Power et al., 2017, 2019) have demonstrated that subjects in resting-state fMRI recording conditions exhibit frequent breathing changes and reflexes (e.g. ‘yawns’ and ‘sighs’). These behaviors likely induce variation in arterial CO2 concentration in the cerebral vasculature, thereby modulating cerebral blood flow. Our findings that the topography and spatiotemporal structure of global fMRI fluctuations is maintained under experimental suppression of PETCO2 fluctuations suggest that global fMRI fluctuations are unlikely to arise primarily from arterial CO2 fluctuations. Another physiological source, high-frequency (aliased) cardiac pulsations, was also determined not to be a significant contributor to fluctuations of the global fMRI signal, as these effects were largely attenuated in ME-ICA denoised time courses (**Supplementary Figure 12**). In addition, the global fMRI signal is still present in high-sampling rate fMRI scans (2.6Hz) with the cardiac cycle spectrally removed through bandpass filtering (free breathing and clamped CO2 dataset; **Supplementary Figure 2**)

The tight coupling between global fMRI and ANS-mediated physiological dynamics observed at rest, and their reproduction by experimental or spontaneous variation in autonomic outflow, provide evidence that the ANS (primarily the sympathetic nervous system) is the key ‘upstream’ driver of low-frequency global fMRI signals. Yet, these results leave open the proximal causal mechanism by which ANS activity and global fMRI signals are linked. Such a link may be direct, via sympathetic-innervation of the cerebral vasculature. Cerebral blood vessels, including the choroid plexus lining the ventricles, are profusely innervated by α-adrenergic receptors that may play a role in cerebral autoregulation (Koep et al., 2022; Lindvall & Owman, 1981). One source of cerebral vasoconstrictive effects may be the superior cervical ganglion that innervates blood vessels in the head and neck (Duyn et al., 2020; Hamel, 2006). Central control of cerebral blood flow from projections of noradrenergic brainstem nuclei (e.g. locus coeruleus) may also have a role (Bekar et al., 2012; Raichle et al., 1975). Currently, the extent of a noradrenergic vasoconstrictive effect (peripheral or central) on cerebral blood vessels, and thereby fMRI signals in the brain, is controversial (van Lieshout & Secher, 2008). Some evidence suggests that sympathetic regulation of cerebral blood flow may be most prominent at frequencies higher than 0.05Hz (Hamner et al., 2010), around the frequency range of fluctuations in global fMRI signals observed in this study. Under this interpretation, systemic vasoconstriction of cerebral arteries/arterioles produces a reduction of cerebral blood volume, inducing a reduction in fMRI signal in the gray matter and white matter (with a delay) and an increase in signal in CSF compartments due to the differential effects of blood volume changes.

The global fMRI signal exhibits a bimodal response to arousal induction (**Figure 2**), and the initial overshoot of this signal (∼4s post onset) is unlikely to be explained by sympathetic modulation of the cerebral vasculature. The temporal alignment of the global fMRI overshoot is within the time to peak (∼4 - 6s) of the canonical hemodynamic response (Buxton et al., 2004). Further, at peaks of the global fMRI signal, early signal increases are prominent in areas of high noradrenergic transport (NAT) density within the sensorimotor cortices (**Figure 3B**; (Ding et al., 2010), suggestive of neurovascular contributions to this area’s signal from noradrenergic projections. Previous research has observed that noradrenergic innervation is characterized by a density gradient that peaks in the sensorimotor cortex and decreases rostrally and caudally (Gaspar et al., 1989).

Large-scale electrophysiological signals from the surface of the brain are commonly observed under abrupt changes in arousal (Cash et al., 2009; Liu et al., 2018; Raut et al., 2021), consistent with our findings of prominent broadband EEG power fluctuations in response to arousal induction (**Figure 2**). The time-to-peak of the global fMRI signal and broadband EEG power were largely similar at around four seconds from stimulus onset. Simultaneous (zero-lag) correlations were also observed in cross-correlations of broadband EEG power with the global fMRI signal under resting-state conditions (**Figure 1**). The similar timing of the global fMRI and broadband EEG response to arousal induction suggests that the electrophysiological potential changes recorded from the EEG are not directly responsible for the fMRI signal fluctuations, as a time-delay would be expected for neurovascular coupling. However, their joint timing in response to arousal may point to a common source in subcortical ascending arousal system nuclei (e.g. locus coeruleus and basal forebrain). Such a hypothesis is supported by evidence that inactivation of these regions (e.g. basal forebrain) leads to suppression of the global fMRI signal (Turchi et al., 2018), and that fMRI signal changes in the basal forebrain occurred together with peaks in the global signal (Liu et al., 2018).

The functional significance of our findings is underscored by the prominence of these low-frequency fluctuations across multiple organ systems. Their induction by arousal suggests that the recruitment of these systems is in service of priming the body for action through heightened sensory receptivity and the shunting of metabolic resources to vital organs, including the brain. The concomitant electrophysiological and hemodynamic (and possibly glymphatic) changes in the brain likely serve the same objective. As one of the most metabolically active organs of the body, the supply of oxygenated blood to the brain is finely regulated to accommodate local increases in metabolism. Brain-wide fluctuations in cerebral blood flow, either arising from systemic vascular (vasoconstriction) or neurovascular mechanisms, may be an anticipatory mechanism of future increases in metabolic rate. Alternatively, it may anticipate or support a physiological process separate from oxygen metabolism, such as neurotransmitter synthesis or regulation of neuronal excitability (Drew, 2022). In this scenario, arousal-induced fluctuations of the global fMRI signal represent a component of a more general recruitment of multiple organ systems to anticipate future action.

Global fluctuations are a prominent component of low-frequency (0.01-0.1Hz) fMRI signal recordings, explaining around 25% of the variance in whole-brain fMRI signals (**Supplementary Figure 2**). At the same time, the majority of variance in the low-frequency fMRI signal is uncorrelated with this component, which may reflect mechanisms separate from those underlying the global fMRI signal. It is likely that certain ‘downstream’ signal dynamics of the global fMRI signal (e.g. vein effects, white matter, CSF) are unrelated to neurovascular coupling mechanisms. However, our findings suggest that global fMRI fluctuations and its concomitant physiological signal fluctuations are a regularly observed feature of the sympathetic-mediated arousal response, and likely plays a crucial role in normal physiological functioning. Taken together, the current findings provide novel insights into the origins and functional significance of global fMRI signals.

## Acknowledgements

BTTY is supported by the NUS Yong Loo Lin School of Medicine (NUHSRO/2020/124/TMR/LOA), the Singapore National Medical Research Council (NMRC) LCG (OFLCG19May-0035), NMRC CTG-IIT (CTGIIT23jan-0001), NMRC STaR (STaR20nov-0003), Singapore Ministry of Health (MOH) Centre Grant (CG21APR1009), the Temasek Foundation (TF2223-IMH-01), and the United States National Institutes of Health (R01MH120080 & R01MH133334). Any opinions, findings and conclusions or recommendations expressed in this material are those of the authors and do not reflect the views of the Singapore NMRC, MOH or Temasek Foundation. CC acknowledges support from NIH grants RF1MH125931 and P50MH109429, and from the Advanced MRI Section of the Intramural Research Program of the NINDS, NIH. LQU is supported by R21HD111805 from NICHD and U01DA050987 from NIDA.

## Supplementary Materials

### Supplementary Movies

Supplementary Movie 1. **Temporal Reconstruction of CPC1 in ME-REST Dataset.**

Supplementary Movie 2. **Temporal Reconstruction of CPC1 in ME-REST-SUPP Dataset.**

Supplementary Movie 3. **Cross-Correlation of fMRI Signals and PPG Amplitude in ME-REST Dataset.**

Supplementary Movie 4. **Cross-Correlation of fMRI Signals and PPG Amplitude in ME-REST-SUPP Dataset.**

### Supplementary Tables

**Supplementary Table 1.** Dataset Details and Demographics. Details (manuscript label, signals recorded, reference) and demographics (sample size, age range, sex) for datasets used in this study. Note, demographics are based on the dataset sample after the quality control stage.

| Dataset | Sample | Physio Signals Used | Source | Sample Size | Number of Sessions | Age Range | Sex (F) |
| --- | --- | --- | --- | --- | --- | --- | --- |
| ME-REST | Full | EEG, Respiration, PPG | Goodale et al. (2021). *ELife*. https://doi.org/10.7554/eLife.62376 | 11 | 15 | 21-35 | 6 |
| ME-TASK | Full | EEG, Respiration, PPG | Unpublished dataset; https://www.changlab.net/ | 6 | 9 | 22-57 | 4 |
| ME-TASK-CUE | Full | EEG, Respiration, PPG | Goodale et al. (2021). *ELife*. https://doi.org/10.7554/eLife.62376 | 12 | 12 | 21-33 | 6 |
| ME-REST-SUPP | Subset | Respiration, PPG | https://openneuro.org/datasets/ds003592/versions/1.0.11<br>**OpenNeuro Accession Number:** ds003592<br>**Version:** 1.0.11 | 87 | 165 | 18 - 34 | 58 |
| HCP-REST | Subset | Respiration, PPG | https://www.humanconnectome.org/ | 30 | 30 | 22-37 | 17 |
| NKI-TASK | Subset | Respiration, PPG, Skin Conductance | http://fcon_1000.projects.nitrc.org/indi/enhanced/ | 50 | 50 | 15-45 | 30 |
| NKI-REST | Subset | Respiration, PPG, Skin Conductance | http://fcon_1000.projects.nitrc.org/indi/enhanced/ | 50 | 50 | 18-45 | 33 |
| NATVIEW-REST | Full | EEG, Pupillometry | (Telesford et al., 2023) | 21 | 33 | 22-51 | 10 |
| YALE-REST | Full | Pupillometry | https://openneuro.org/datasets/ds003673/versions/2.0.1<br>**OpenNeuro Accession Number:** ds003673<br>**Version:** 2.0.1 | 27 | 54 | 21-37 | 16 |
| Clamped CO2 | Full | N/A | (Golestani & Chen, 2020) | 13 | 13 | 18-32 | 9 |

### Supplementary Figures

**Supplementary Figure 1.**
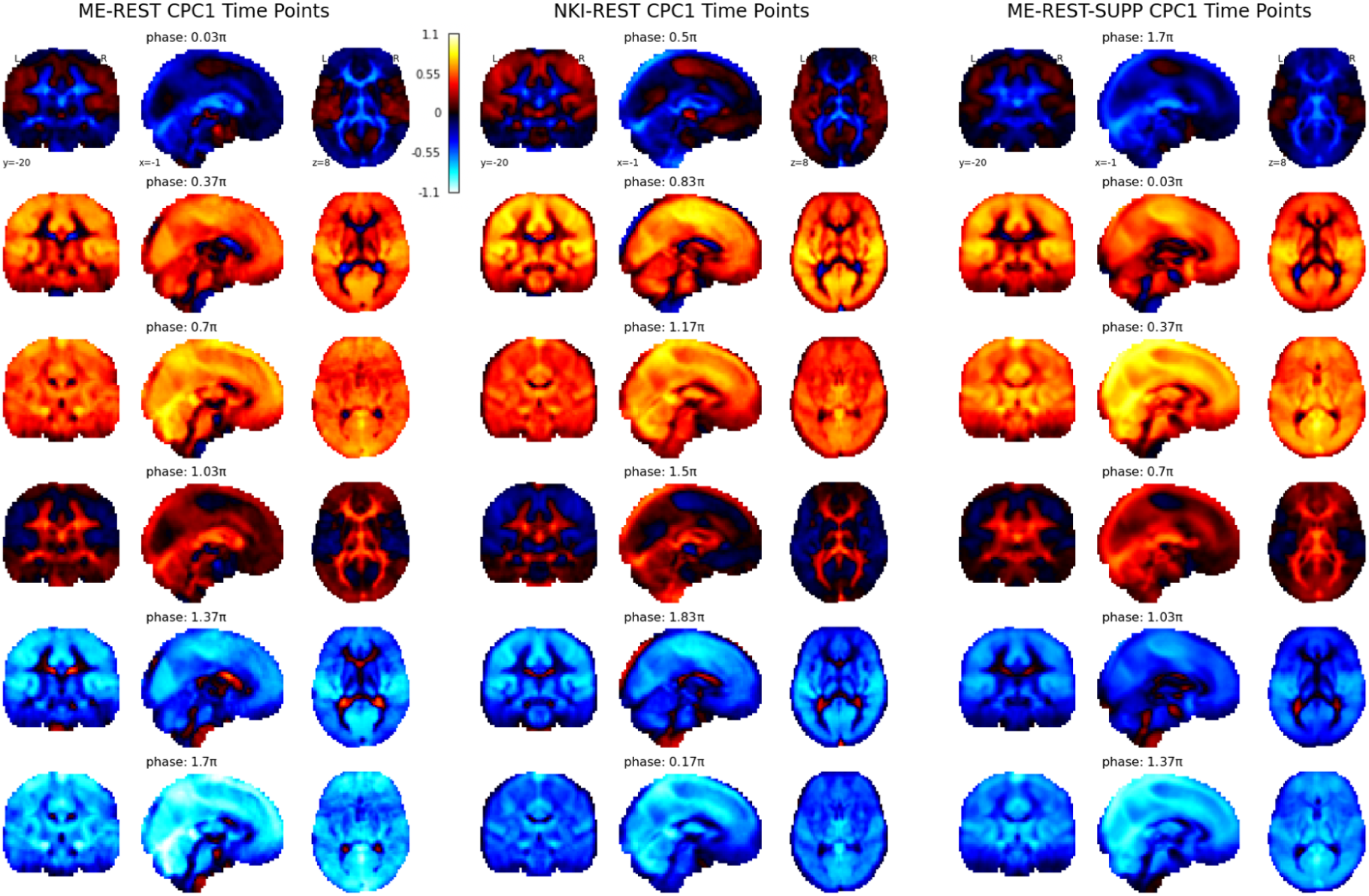
Spatiotemporal Dynamics of Global fMRI Signals. Temporal reconstruction of the first complex principal component (CPC1) of three resting-state fMRI datasets (ME-REST, NKI-REST, ME-REST-SUPP). Complex PCA (CPCA) on other resting-state datasets (HCP-REST, NATVIEW-REST, YALE-REST) yielded similar results, and are not included here for space. As shown in previous work, the first principal component from PCA and complex principal component from CPCA extract the global fMRI signal, with CPCA yielding a spatiotemporal representation of the global fMRI signal (Bolt et al., 2022). The time course of a complex principal component is represented in complex numbers, and its phase can be extracted (measured in radians). A temporal reconstruction of the complex principal component can be constructed via averaging of the original fMRI signals (in voxel space) at similar phase values. We selected six equally spaced phase values to display the spatiotemporal dynamics of the global fMRI signal. Time moves in the positive direction, such that increasing phase values move forward in time. A consistent spatiotemporal pattern is observed across datasets: a global increase in fMRI signals in the gray and white matter followed by a propagation of fMRI signals to large draining veins and ventricles, and then a global decrease in fMRI signals.

**Supplementary Figure 2.**
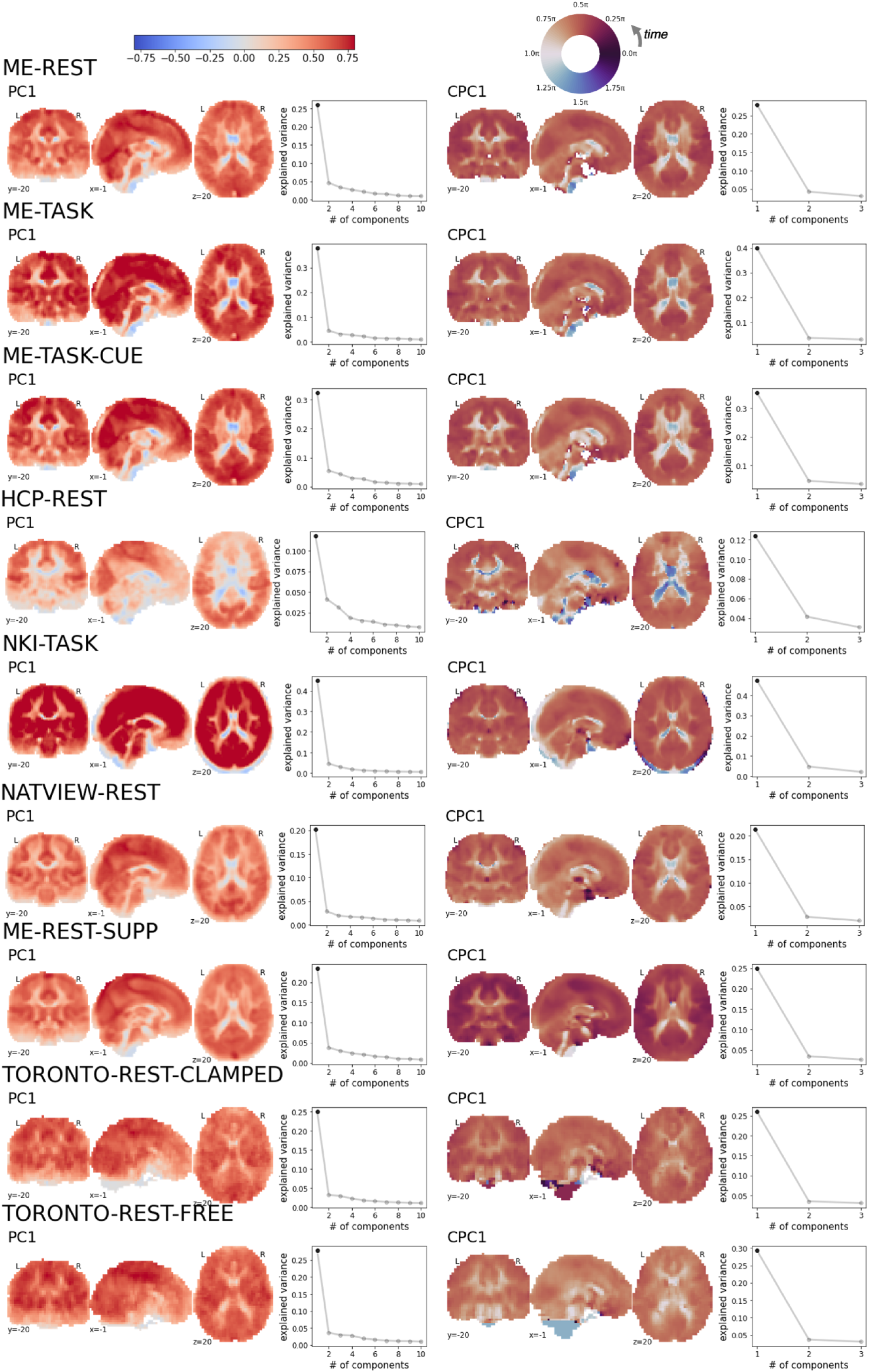
First Principal Component and Complex Principal Component Across Datasets. Spatial weights of the first principal component (PC1; left) and phase delay maps of the first complex principal component (CPC1; right) across all datasets used in this study. Explained variance plots (Scree plots) are displayed to the right of each brain map displaying the explained variance by the first and subsequent principal components. The phase delay map of the first complex principal component encodes the time-delay (in radians) between voxels within the component. Because phase delay is measured in radians (0 to 2pi), they are displayed with a circular color map.

**Supplementary Figure 3.**
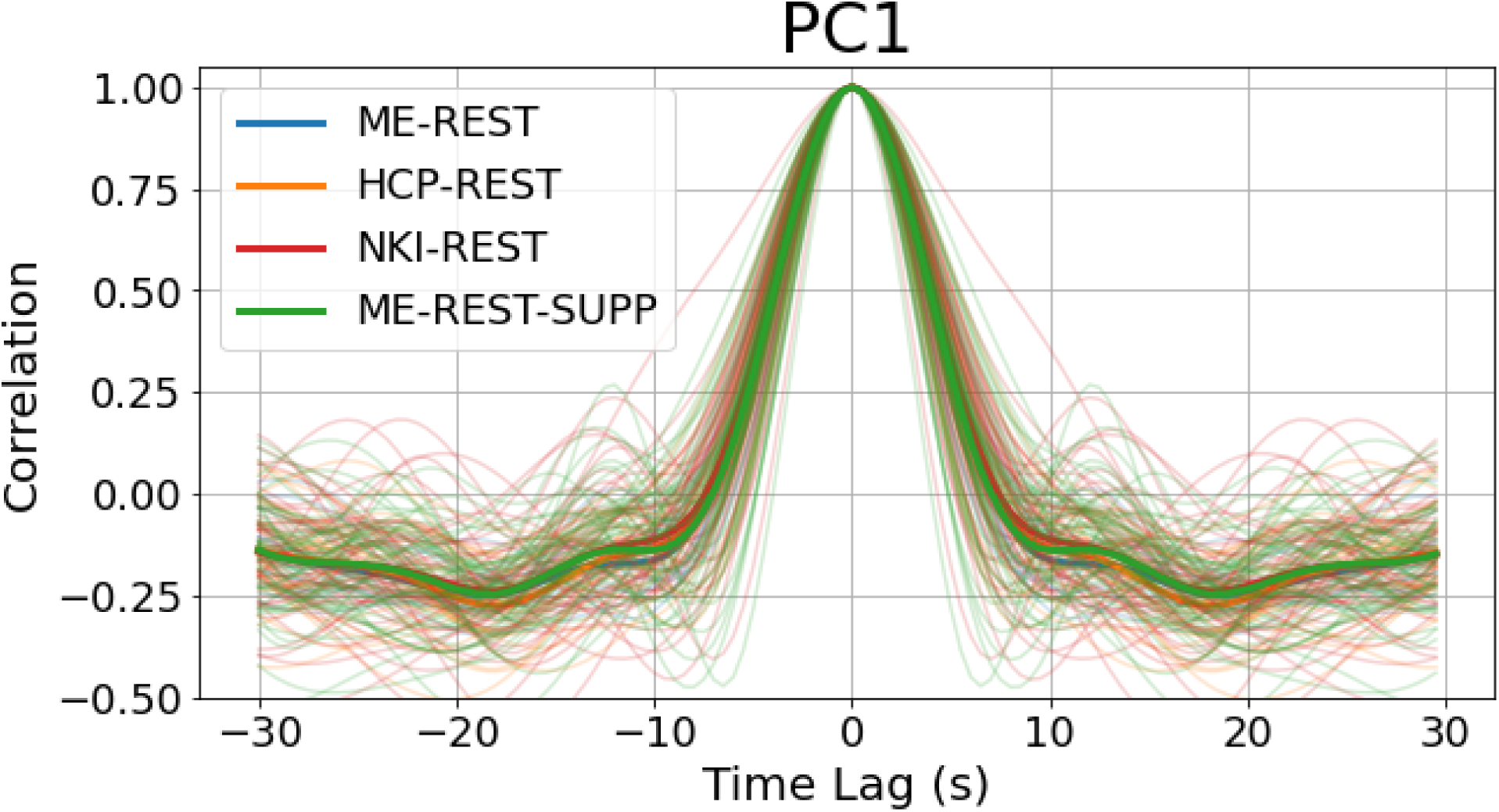
Auto-Correlation Plot of PC1 Time Courses. The cross-correlation of the PC1 time course with itself from the ME-REST, HCP-REST, NKI-REST and ME-REST-SUPP datasets. Subject-level auto-correlations are displayed in lighter colors, and the group-average auto-correlation is displayed with a thicker, darker line.

**Supplementary Figure 4.**
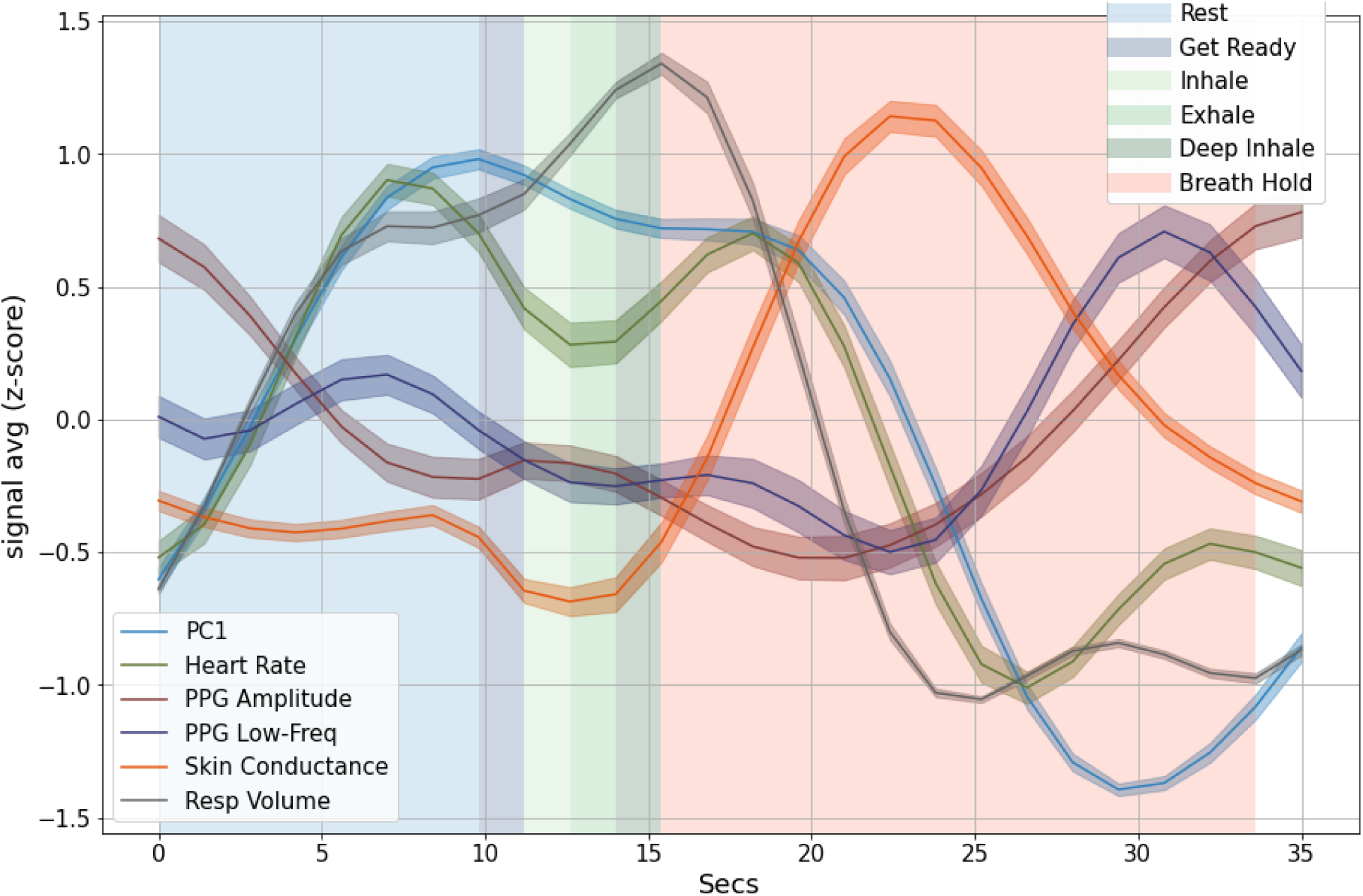
NKI-Breath Hold Task. Physiological trial averages for a full block of the paced inhalation/breath hold task (NKI-TASK). The paced breathing/breath-hold task was a block design consisting of a fixed sequence of rest (10s), a cue (2s) followed by two deep inhalations (6s), and a breath-hold (18s), immediately followed by another sequence. Time points in separate blocks are shaded to distinguish activity in each block: rest (10s duration; blue), ‘get ready’ cue (2s; dark blue), inhalation (2s; light green), exhalation (2s; medium green), deep inhalation (2s; dark green) and breath hold (18s; red).The physiological dynamics of the paced breathing/breath hold task (NKI-TASK) are more complex than those observed in response to isolated deep inhalations (Figure 2A). For example, large increases in respiratory volume are observed during the paced inhalation exercise before the breath hold and the ‘rest’ blocks that precede the paced inhalation exercise, due to their placement immediately after the breath hold. The timing of the large amplitude peak of global fMRI signals in the rest period (∼10s) and following the deep inhalation block (19s) are consistent with the timing of the respiration response peak observed to isolated deep inhalations (Figure 2A). Consistent with the response to isolated deep inhalations, an increase in heart rate is observed around the time of the global fMRI peak, shortly followed by peripheral vasoconstriction. In addition, a large amplitude response in skin conductance is observed around the same time as peripheral vasoconstriction. The trough of the global fMRI response, along with peripheral vasodilation, occurs in the latter half of the breath hold block, consistent with the peak-to-trough timing observed in deep inhalations.

**Supplementary Figure 5.**
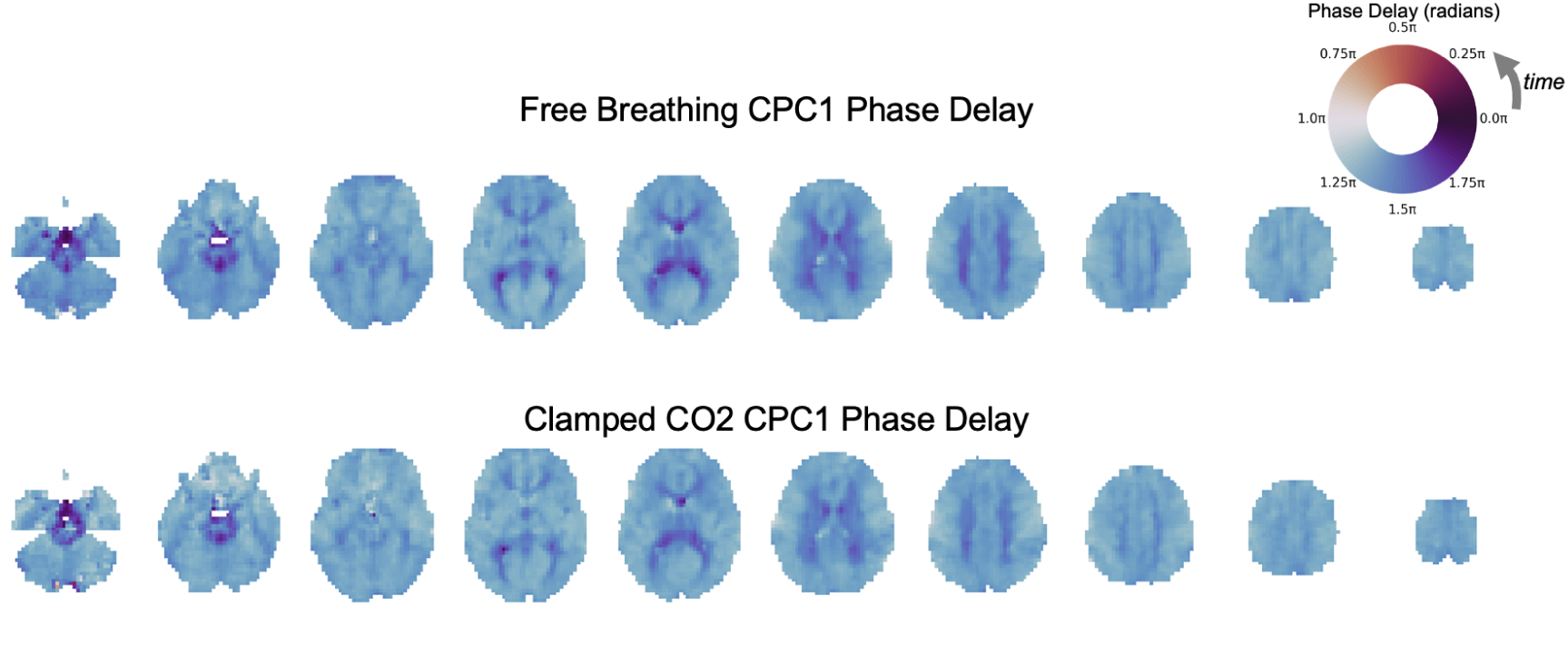
First Complex Principal Component Phase Delay in Free Breathing and Clamped CO2 Conditions. Phase delay maps of the first complex principal component (CPC1) computed from fMRI time courses in free breathing and clamped CO2 conditions. The phase delay map of the first complex principal component encodes the time-delay (in radians) between voxels within the component. Because phase delay is measured in radians (0 to 2pi), they are displayed with a circular color map. As can be observed from the brain maps, the distribution of phase delay values across the brain is highly similar across free breathing conditions and clamped conditions.

**Supplementary Figure 6.**
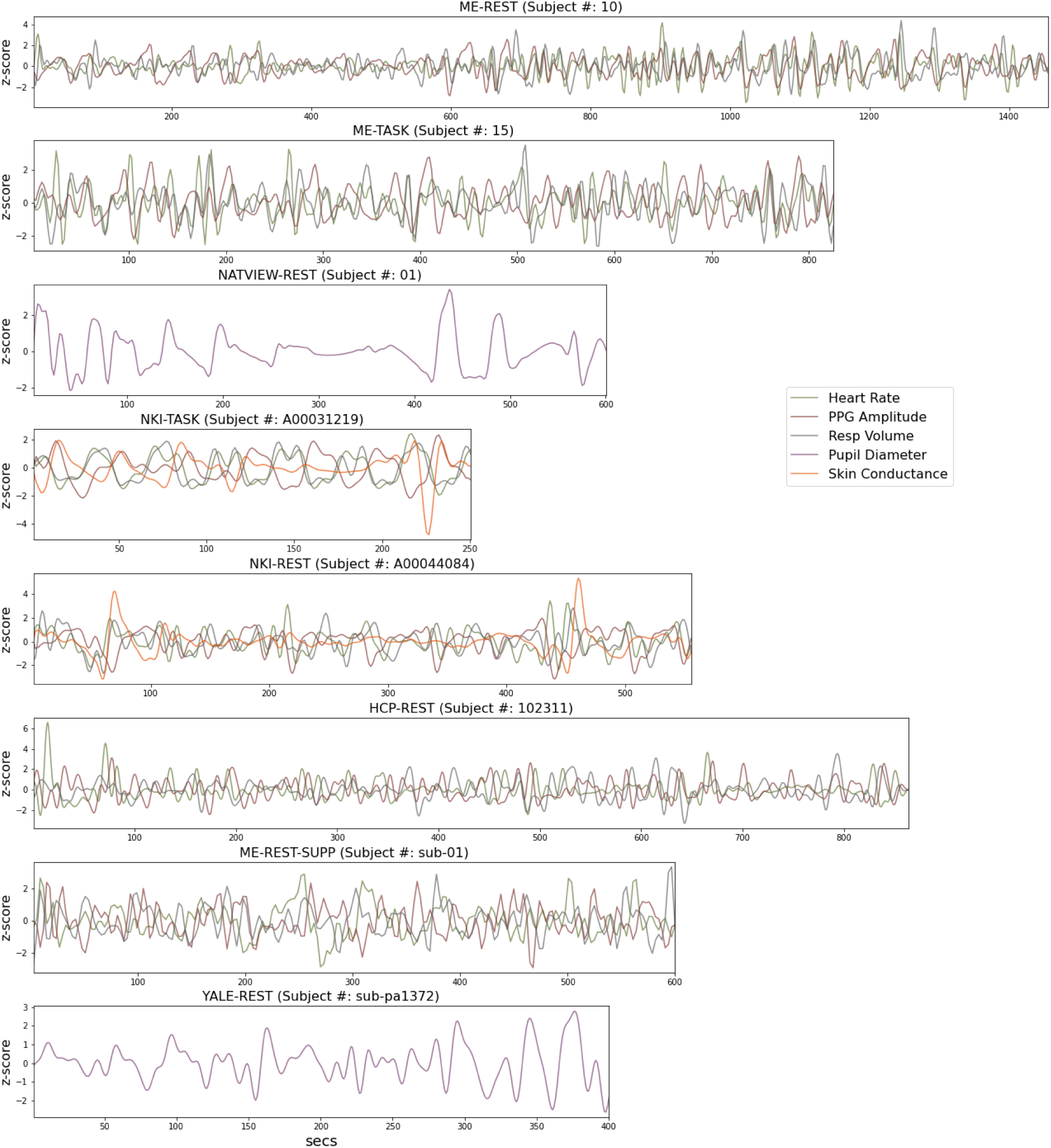
Physiological Time Series from Example Subjects.

**Supplementary Figure 7.**
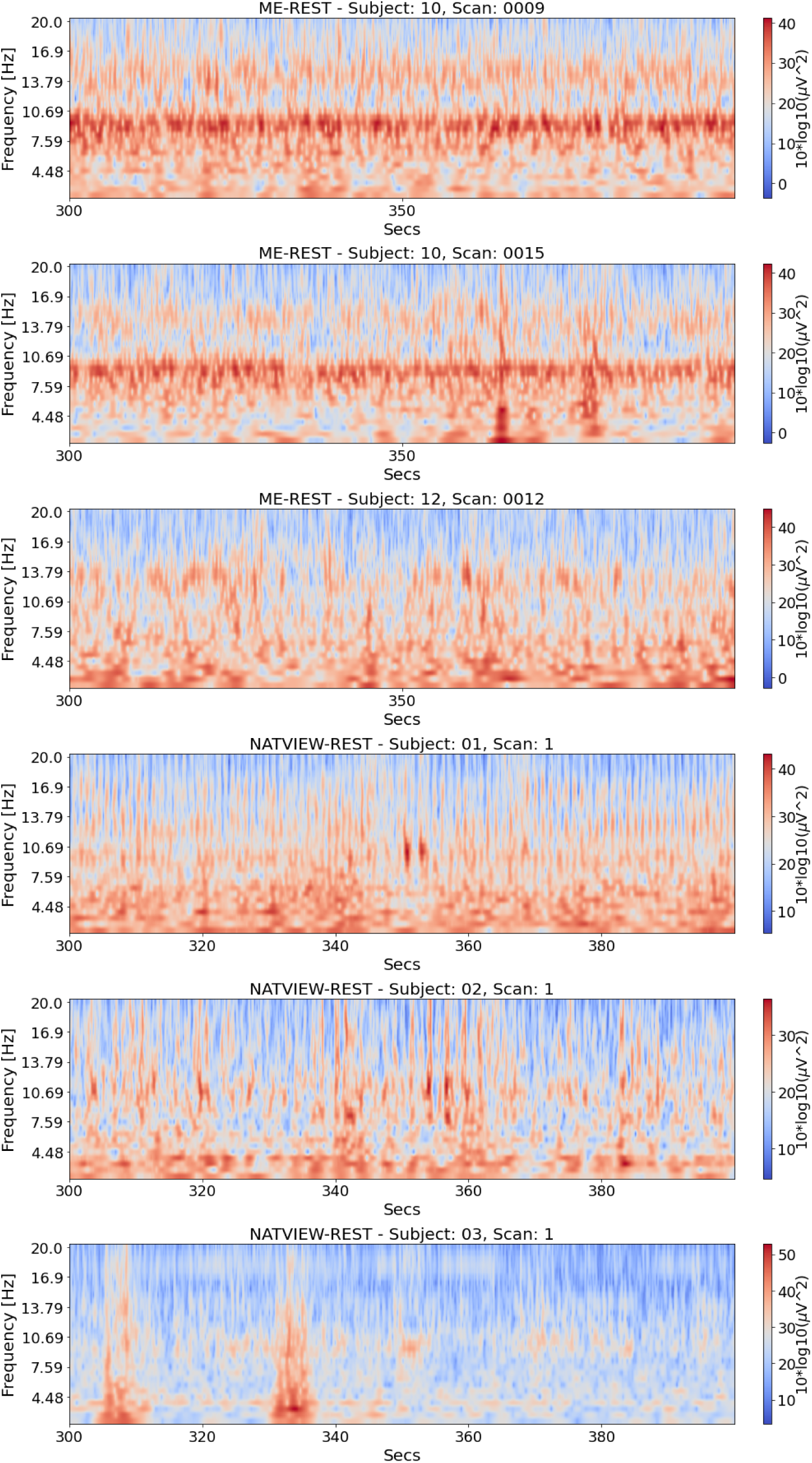
Channel-Averaged EEG Spectrograms Samples from Example Subjects. EEG time-frequency power plots of sample time points (300 - 400 seconds post scan-onset) averaged across channels used in the study (posterior and occipital channels). Time-frequency EEG power was extracted via Morlet wavelet filters using the same parameters for analyses presented in Figure 1 and Figure 2 (number of cycles = 15; frequencies: 2 - 20Hz).

**Supplementary Figure 8.**
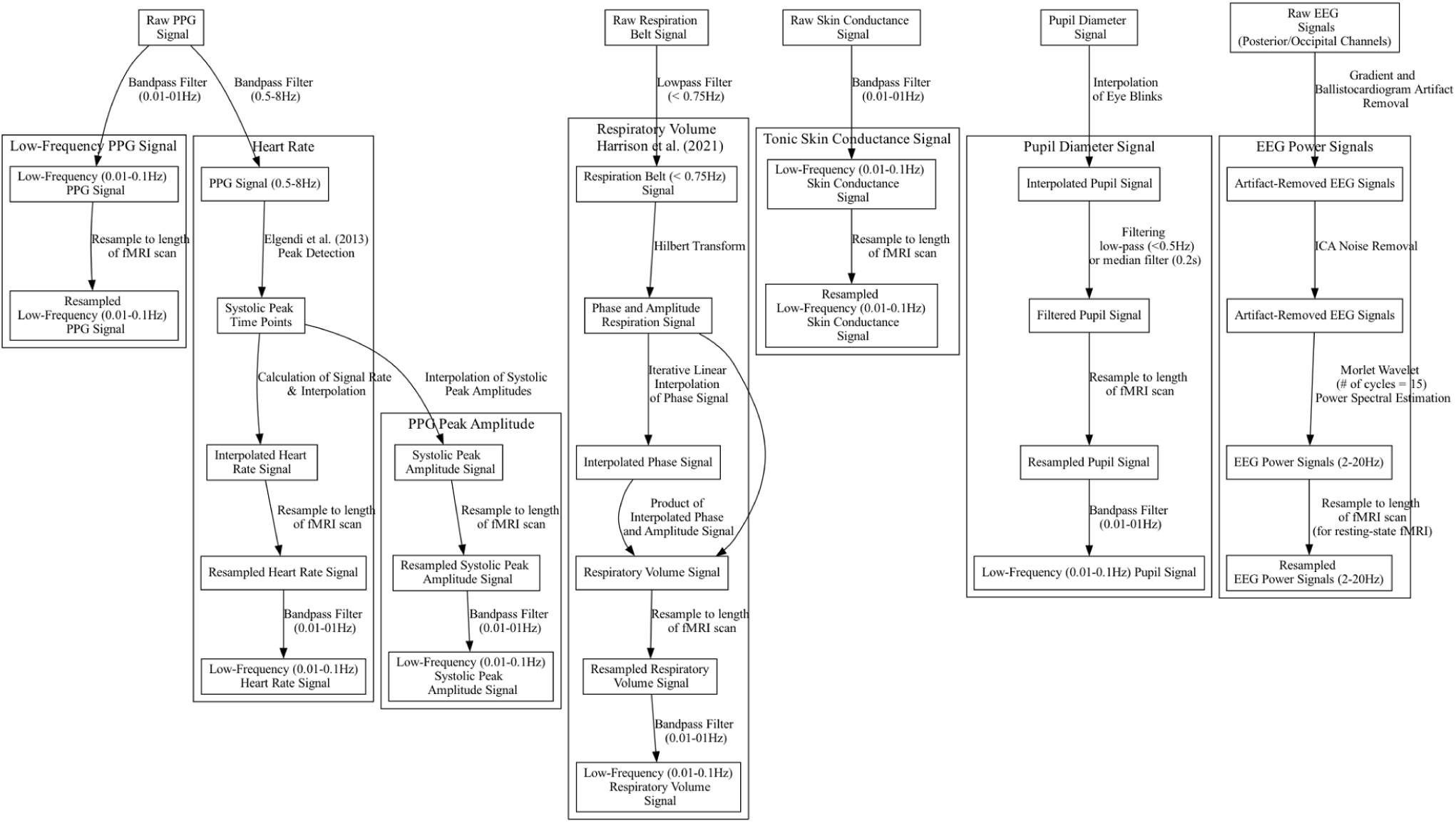
Electrophysiological Signal Preprocessing Pipeline.

**Supplementary Figure 9.**
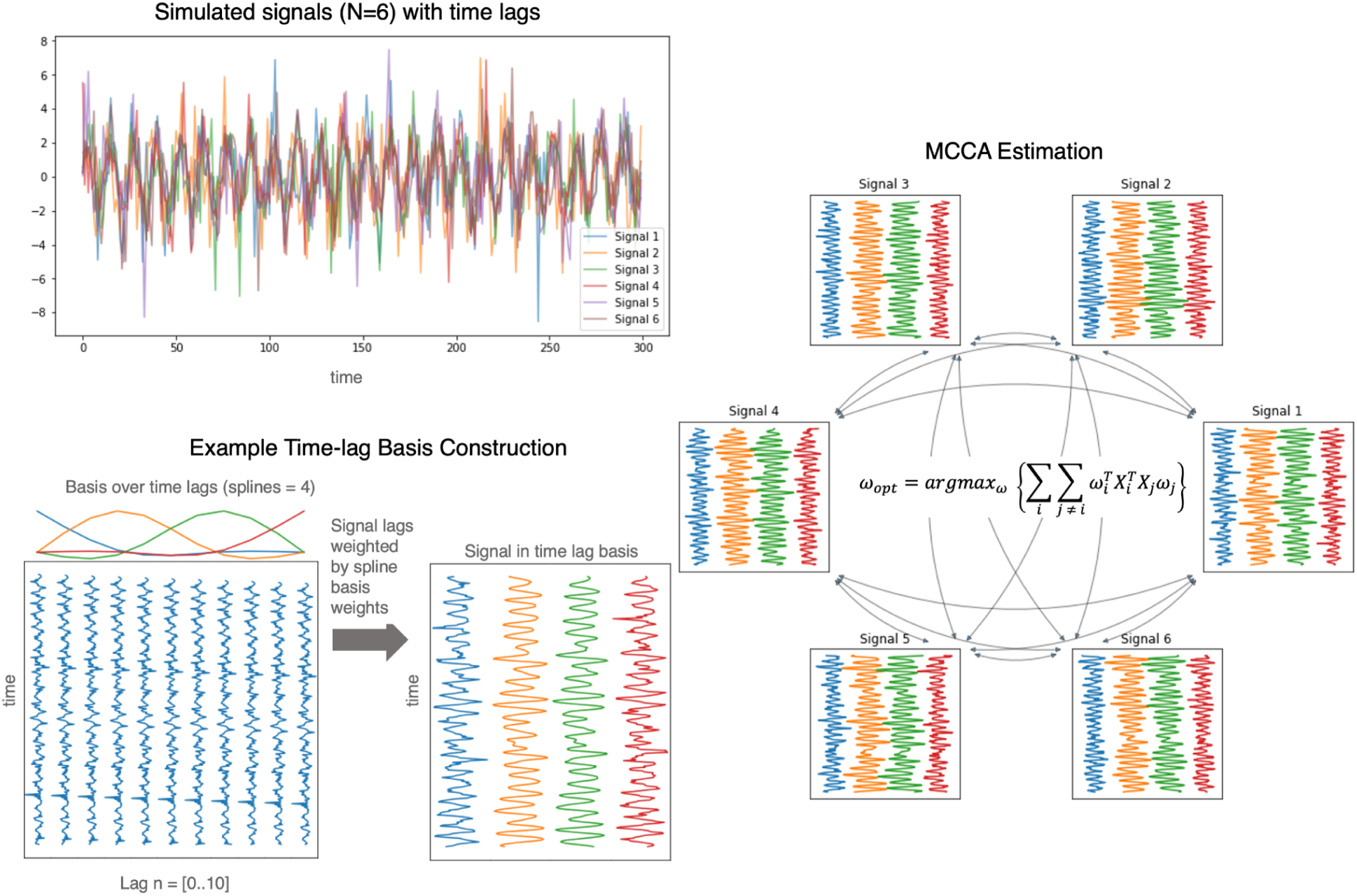
Illustration of Multiset Canonical Correlation Analysis.

**Supplementary Figure 10.**
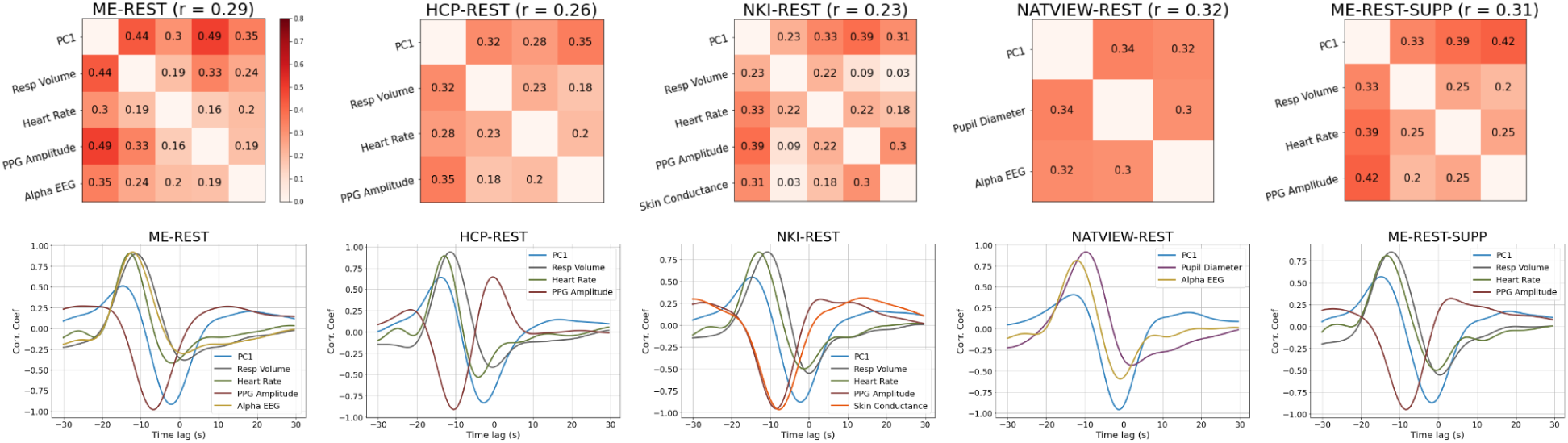
MCCA of Resting-State Datasets and Cross-Correlations. **Top panel:** The pairwise correlations (top) between all physiological signals (including the global fMRI signal; PC1) in the first canonical component of the MCCA analysis and their time lags (bottom) for five resting-state datasets, as displayed in Figure 1. The average pairwise correlation is displayed beside the title of each correlation matrix. **Bottom panel:** To extract timing information between the signals in this low-dimensional space, we cross-correlated each physiological signal with its projection onto the first canonical component. The cross-correlations between each physiological signal and its projection onto the first canonical component are displayed in the bottom panel, where each signal is displayed in a different color. Comparison of the relative timing between peaks of the cross-correlation curves across physiological signals provides the lead-lag relationships between signals within the first canonical component of MCCA.

**Supplementary Figure 11.**
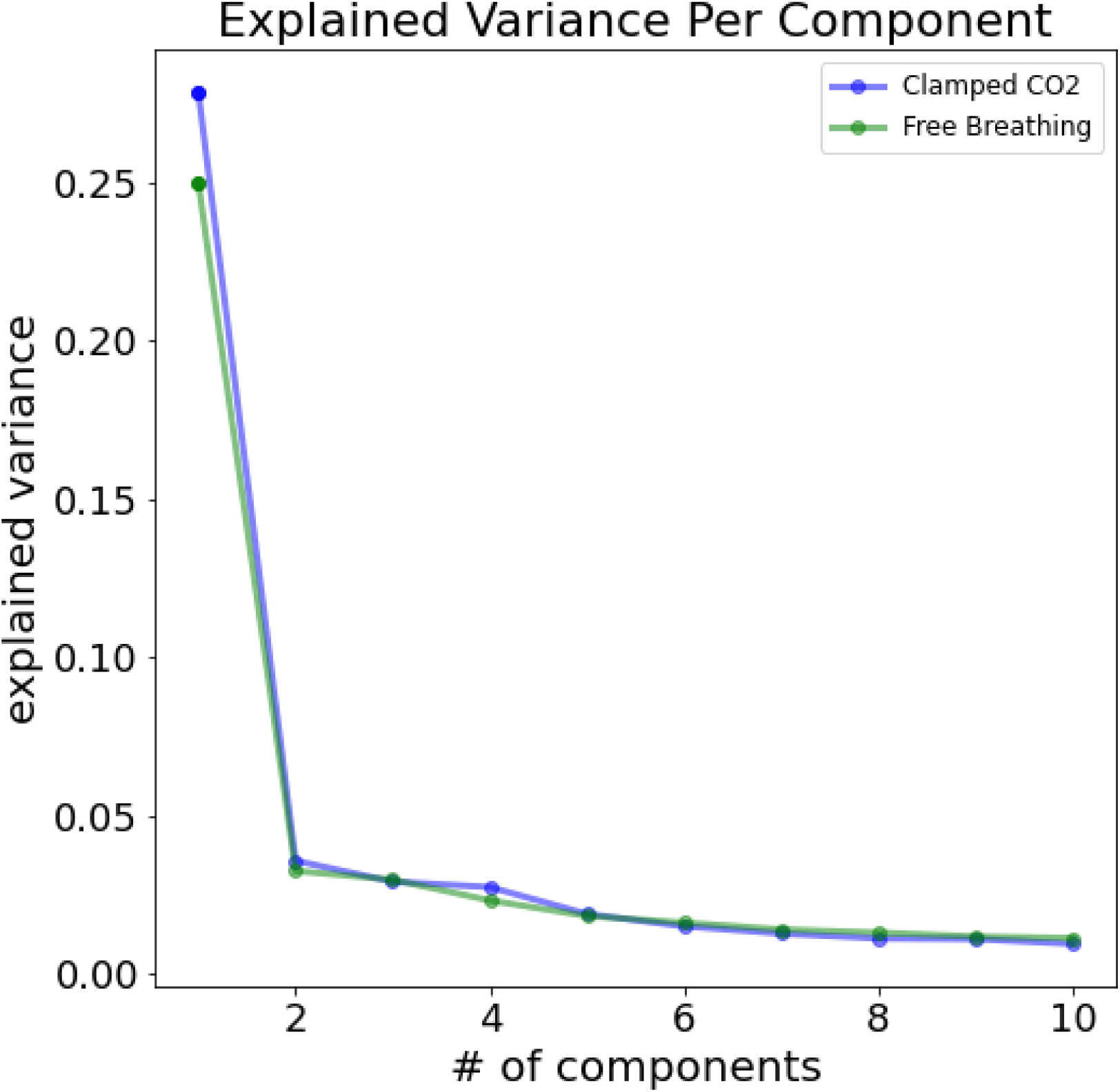
Scree Plot of Eigenvalues for Clamped CO2 and Free Breathing Conditions. A scree plot of eigenvalues of the first ten components from PCA estimated from the free breathing (green) and ‘clamped’ CO2 (blue) condition. As can be observed, the spatial distribution of the global fMRI signal is maintained when variations of CO2 are experimentally suppressed. More generally, the low-dimensional spatial structure of fMRI time courses between the two conditions, as reflected in the scree plot, is similar between the two conditions.

**Supplementary Figure 12.**
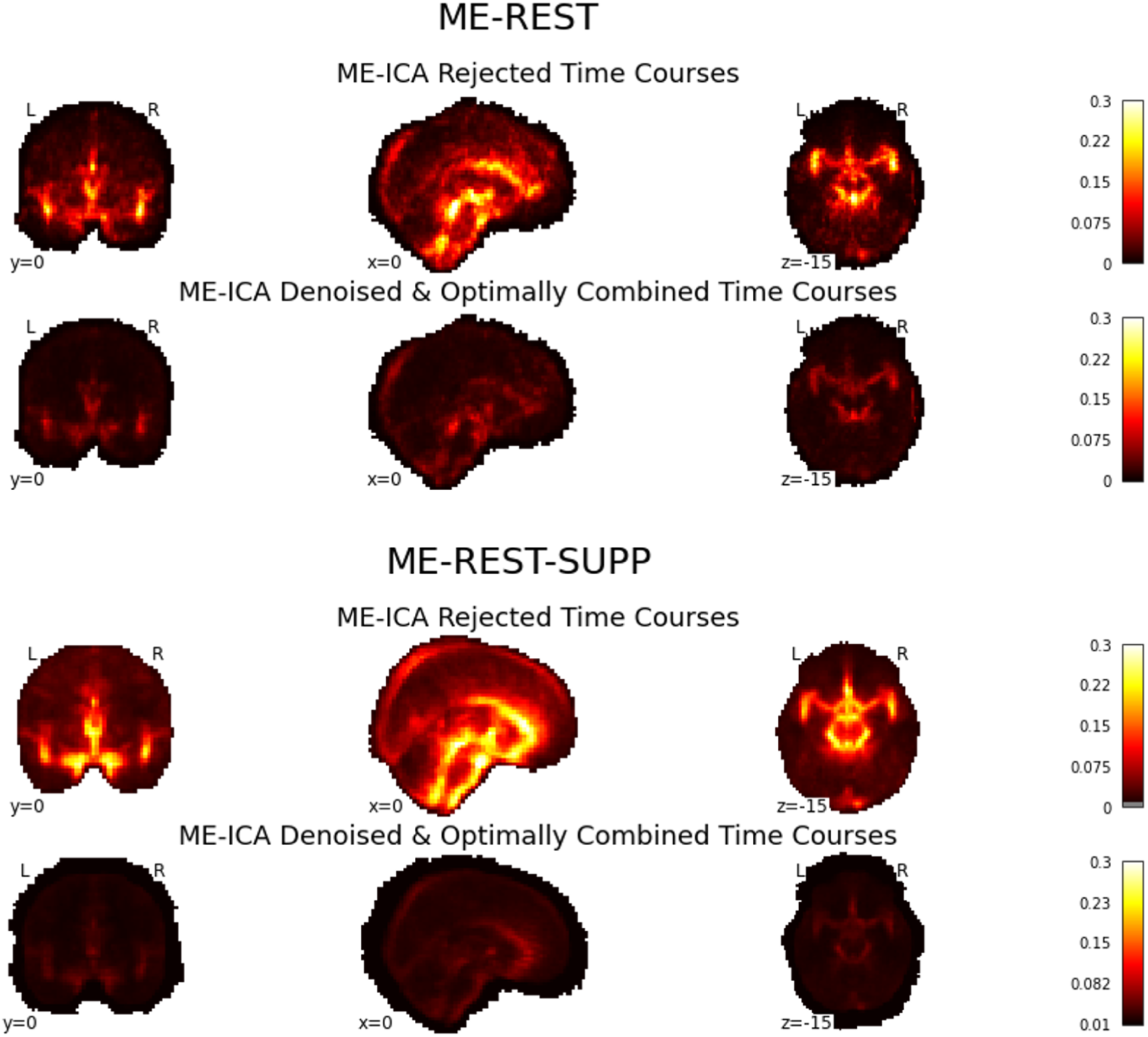
Explained Variance of High-Frequency Cardiac Regressors In Multi-Echo ICA Denoised Time Courses. Brain maps of the group-averaged explained variance (R^2^) estimates at each voxel by high-frequency cardiac (RETROICOR) regressors. To verify that the spatiotemporal pattern of the global fMRI signal (Figure 3) does not arise primarily from aliased cardiac pulsations, we performed regression modeling of (aliased) high-frequency cardiac RETROICOR regressors in Multi-Echo (ME) fMRI data. Specifically, we performed regression modeling on both the ME-ICA denoised time courses and on the time courses that were rejected as echo-time independent ‘noise’ by ME-ICA (i.e., time courses constructed from the rejected ME-ICA components). As can observed in the group-averaged R^2^ maps, high-frequency cardiac pulsation effects are primarily observed in the ME-ICA rejected time courses, and are largely attenuated in the denoised time courses.

### Methods and Materials

#### Participants and Data Acquisition

Ten datasets were analyzed in this study: 1) a simultaneous EEG/multi-echo resting-state fMRI dataset (ME-REST), 2) a simultaneous EEG/multi-echo respiration task fMRI dataset (ME-TASK), 3) a simultaneous EEG/multi-echo reaction time task fMRI dataset (ME-TASK-CUE) 4) a multiband accelerated single-echo resting-state fMRI dataset from the Human Connectome Project (Van Essen et al., 2013) (HCP-REST), 5) a multiband accelerated single-echo fMRI dataset from the enhanced Nathan Kline Institute (NKI) - Rockland Sample (Nooner et al., 2012), including a resting-state (NKI-REST) and 6) respiration task (NKI-TASK), 7) a supplementary multi-echo fMRI resting-state dataset (Spreng et al., 2022)

(ME-REST-SUPP), 8) a multiband accelerated single-echo resting-state fMRI dataset with simultaneous pupillometry from Yale University (Lee et al., 2022) (YALE-REST), 9) a simultaneous EEG/single-echo resting-state fMRI dataset from NKI (Telesford et al., 2023) (NATVIEW-REST), 10) a multiband accelerated single-echo resting-state fMRI dataset recorded under a free breathing condition and a PETCO2 condition clamped to the average PETCO2 for each participant. Dataset details and demographics are presented in **Supplementary Table 1**.

#### ME-REST, ME-TASK, ME-TASK-CUE Data

Simultaneous multi-echo fMRI-EEG eyes-closed resting-state scans (ME-REST) were acquired from 11 healthy, right-handed participants (6 females, mean age = 25.9 years). All subjects provided written informed consent, and human subjects protocols were approved by the Institutional Review Boards of the National Institutes of Health and Vanderbilt University. Two resting-state sessions were recorded for four of the subjects, creating a total of 15 scans. Simultaneous multi-echo fMRI-EEG respiration task scans (ME-TASK) were acquired under the same acquisition protocol for six healthy, right-handed participants (4 females, mean age = 30.5). Two task sessions were recorded for three of the subjects, creating a total of 9 scans.

The respiration task experimental design consisted of a sparse event-related design with instructions to the participants to take a deep breath in response to an auditory cue (a constant tone). The interstimulus interval (ISI) between auditory cues was randomly jittered between the range of 60.55 and 131.25 sec. Auditory cue timing was consistent across scans. One participant overlapped between the resting-state and respiration task sessions. Simultaneous multi-echo fMRI-EEG reaction time task scans (ME-TASK-CUE) were acquired under the same acquisition protocol for twelve healthy, right-handed participants (6 females, mean age = 25.6 years). The reaction time task experimental design consisted of a sparse event-related design with instructions to the participants to press a button in response to an auditory cue. For some subjects (N=5), the ISI between auditory cues was set to 35s±6s, and for the rest (N=7) the ISI was randomly jittered between 8 and 89 secs. Nine participants overlapped between the resting-state and reaction time task sessions.

Detailed MRI/EEG acquisition parameters are provided in Goodale et al. (2021). Briefly, anatomical T1-weighted structural and multi-echo EPI functional scans were collected on a 3T Siemens Prisma scanner with a Siemens 64-channel head/neck coil. The multi-echo EPI sequence was acquired with TR = 2100 ms, echo times = 13.0, 29.4, and 45.7 ms, flip angle = 75 degrees, and voxel size = 3mm isotropic. The duration of the resting-state scan was 24.5 minutes, corresponding to a total of 700 volumes. The duration of the respiration task scan was slightly variable (14-15 minutes across subjects), corresponding to a total of 400-435 volumes, depending on the subject. Simultaneous scalp EEG (sampling rate = 5 kHz) was acquired during the resting-state and respiration task using a 32-channel MR-compatible system.

Photoplethysmography (PPG) and respiration belt signals (sampling rate: 2 kHz) were acquired during both the resting-state and respiration task sessions. The PPG transducer was placed on the left index finger. MRI scan triggers were recorded along with EEG and physiological signals for data synchronization.

#### HCP-REST Data

We analyzed eyes-open resting-state fMRI EPI functional scans from the HCP S1200 release. We randomly selected 30 unrelated, healthy young adults (ages 22–37; 17 females) for our study with high quality physiological recordings (confirmed through visual inspection; see note below on quality control procedure). We chose 30 subjects for two reasons: 1) analyses were found to be replicable at small sample sizes (< 15 subjects) for the HCP dataset, and 2) the length of the scans (length of scan: 1200 time points) imposed significant computational challenges. Detailed MRI/Physio acquisition parameters are provided in Smith et al. (2013).

Briefly, resting-state fMRI scans were collected on Siemens 3T Tim Trio scanners with a multiband (factor of 8) accelerated EPI sequence with the following parameters: TR: 720ms, TE = 33.1ms, flip angle = 52 degrees, and voxel size= 2mm isotropic. Resting-state fMRI data was collected over two consecutive days for each subject and two sessions, each consisting of two 15-minute runs, amounting to four resting-state sessions per subject. Within a session, the two runs were acquired with opposite phase encoding directions: L/R encoding and R/L encoding. A single 15 min scan from each participant on the first day of scanning was selected. We balanced the number of L/R and R/L phase encoding scans across our participants (n=15 for L/R direction) to ensure results were not biased by acquisition from any given phase encoding direction. Photoplethysmography (PPG) and respiration belt signals (sampling rate = 400 Hz) were simultaneously acquired with resting-state EPI scans along with MRI scan triggers for data synchronization.

#### NKI-REST and NKI-TASK Data

We analyzed resting-state and task-fMRI breath-hold EPI functional scans, and high-resolution anatomical T1w images from the enhanced Nathan Kline Institute (NKI) Rockland sample. We randomly selected 50 adult subjects with high-quality physiological recordings (confirmed through visual inspection; see note below on quality control procedure) independently for each dataset (NKI-REST: ages 18-45, 33 females; NKI-TASK: ages 15-45, 30 females). Twenty subjects overlapped between the two datasets. A single respiration task session was recorded per participant with a 1400ms TR. Two multi-band accelerated resting-state sessions (TR: 1400ms and TR: 650ms) were collected per participant. We selected the 1400ms (TR) acquisition for consistency with the respiration task. Detailed acquisition parameters are provided on the Enhanced NKI-Rockland webpage (http://fcon_1000.projects.nitrc.org/indi/enhanced/index.html). Briefly, resting-state and breath hold fMRI scans were collected on a Siemens 3T Tim Trio scanner with a multiband (factor of 4) accelerated EPI sequence with the following parameters: TR = 1400ms, TE = 30ms, flip angle = 65 degrees, voxel size = 2mm isotropic. The duration of the resting-state scan was 10 minutes. The respiration task was a block design with the following sequence: 1) a 10-sec rest, 2) a 2-sec visual stimulus indicating the start of the trial (text: ‘Get Ready’), 3) a 2-sec inhalation, 4) a 2-sec expiration, 5) a 2-sec deep inhalation, and 5) a breath hold for 18 secs. This sequence was repeated seven times, for a total duration of 4.5 minutes. PPG signals (recorded from the tip of the index finger), skin conductance (galvanic skin response; recorded from the hand), and respiration belt signals were simultaneously acquired with EPI scans along with MRI scan triggers for data synchronization.

#### ME-REST-SUPP Data

As a supplementary multi-echo fMRI dataset to confirm findings from the ME-REST and HCP-REST datasets, we analyzed eyes-open resting-state data from the neurocognitive aging data release (Spreng et al., 2022). We selected 87 young adults with high quality physiological recordings (confirmed through visual inspection; see note below on quality control procedure) (ages 20 - 34, 20 females). Each participant performed two resting state fMRI sessions during the study visit. Detailed MRI acquisition parameters are provided in Spreng et al. (2022). Briefly, anatomical T1-weighted structural and multi-echo EPI functional scans were collected on a 3T GE Discovery MR750 scanner with a 32-channel head coil. The multi-echo EPI sequence was acquired with TR = 3000 ms, echo times = 13.7, 30, and 47 ms, flip angle = 83 degrees, and voxel size = 3mm isotropic. The duration of the resting-state scan was 10 minutes, corresponding to a total of 204 volumes. Photoplethysmography (PPG) and respiration belt signals (sampling rate: 40 or 50 Hz) were acquired during the session.

#### YALE-REST Data

For analysis of pupillometry signals, we analyzed eyes-open resting-state fMRI data from the Yale Resting-State Pupillometry/fMRI dataset (Lee et al., 2022) via OpenNeuro (Markiewicz et al., 2021; OpenNeuro Accession Number: ds003673). 24 of the 27 participants were selected for analysis based on quality control of the pupil recordings (ages 21 - 37, 16 females). Both resting-state fMRI sessions collected during the study visit were used. Detailed MRI acquisition parameters are provided in Markiewicz et al. (2021). Briefly, anatomical T1-weighted structural and single-echo, multiband EPI functional scans were collected on a MAGNETOM Prisma MRI scanner. The single-echo, multiband EPI sequence was acquired with TR = 1000ms, TE = 30ms, multiband acceleration factor = 5, flip angle = 55 degrees and voxel size = 2mm. The duration of each resting-state scan was 6 minutes and 50s. Simultaneous eye-tracking and pupil dilation was recorded using a MR-compatible infrared EyeLink 1000 Plus eye-tracking system with a 1000Hz sampling rate. We used minimally preprocessed pupil data provided by the authors for analysis (see details below).

#### NATVIEW-REST Data

To replicate EEG and pupillometry associations, we analyzed eyes-open resting-state data with simultaneous EEG-fMRI-pupillometry recordings from NKI (Telesford et al., 2023). We selected 21 of 22 total subjects based on visual inspection of the minimally processed pupil diameter recordings (ages 22-51, 10 females). The resting-state scans were approximately 10 minutes in duration (288 volumes) and were collected as part of a larger naturalistic viewing experiment with several visual and movie watching scans. For some subjects (N=12), two resting-state sessions were collected, and both were used in our analysis. Detailed MRI/EEG acquisition parameters are provided in Telesford et al. (2023). Briefly, anatomical T1-weighted structural and single-echo EPI functional scans were collected on a 3T Siemens TIM Trio scanner with a 12-channel head coil. Resting-state EPI sequences were acquired with TR=2100ms, TE=2500ms, flip angle = 60 degrees, voxel size= ∼3.5mm isotropic.

Simultaneous scalp EEG (sampling rate = 5 kHz) was collected using a customized 61-channel MR-compatible cap, including two electrooculogram (EOG) channels (above and below the left eye) and one electrocardiography (ECG) channel placed on the back. As with the *YALE-REST* dataset, simultaneous eye-tracking and pupil dilation was recorded using an MR-compatible infrared EyeLink 1000 Plus eye-tracking system with a 1000Hz sampling rate. We used minimally preprocessed pupil data provided by the authors for analysis (see details below). Respiratory time courses were also collected with a respiratory belt, but due to data quality issues were not included in our analysis.

#### Free Breathing and ‘Clamped’ CO2 Data

To determine the effect of spontaneous variation in arterial CO2 produced by changes in respiratory depth/rate on global fMRI dynamics, we analyzed resting-state fMRI data from 13 participants (ages 18 - 32, 9 females) collected during free breathing and clamped CO2 conditions (Golestani & Chen, 2020). In the ‘clamped’ condition, participant’s end-tidal CO2 (PETCO2) levels were clamped to their average PETCO2 level using the RespirAct™ breathing circuit (Thornhill Research, Toronto, Canada). For the free-breathing condition, participants were allowed to breathe freely over the course of the scan. Detailed MRI acquisition parameters are provided in Golestani & Chen (2020). Briefly, anatomical T1-weighted structural and single-echo, multiband EPI functional scans were collected on a Siemens TIM Trio 3T MRI scanner with a 32-channel head coil. Free-breathing and clamped CO2 fMRI recordings were acquired with TR = 380ms, TE = 30ms, multiband acceleration factor = 3, flip angle = 40 degrees and voxel size = 4x4x5 mm3.

#### Quality Control of Physiological Recordings

Peripheral physiological recordings tend to be noisy, with relative noise levels dependent on their placement location, subject compliance and movement. To ensure our analysis was not affected by acquisition artifacts of the physiological recordings, raw signals from all subjects in each dataset were visually inspected before inclusion. Several qualitative criteria for each signal type were employed to determine physiological signals for inclusion in the analysis. PPG signals were inspected for 1) clearly visible pulsatile waveforms across the entire recording session and 2) the infrequent occurrence of high amplitude noise artifacts. Respiratory belt signals were inspected for 1) clearly visible respiratory time courses reflecting the peak and trough of the abdomen during the respiratory cycle and 2) the absence of prominent ceiling or floor effects/cut-offs at peaks of the respiratory waveform. The primary criteria for the minimally preprocessed pupil diameter time courses in the NATVIEW-REST dataset were the absence of multiple, long duration (subjectively determined) eye closures. For the pupil diameter time courses in the YALE-REST dataset (Lee et al., 2022), a previous exclusion criteria was applied such that the percentage of missing time points due to eye-closures and blinks did not exceed 35%. Skin conductance time courses for the NKI-REST and NKI-TASK dataset were inspected for 1) prominent low-frequency fluctuations visible from background noise, and 2) the absence of extreme, sudden jumps/drops in baseline signal.

#### MRI Data Preprocessing

Excluding the ME-REST, ME-TASK, ME-TASK-CUE and HCP-REST datasets, datasets were received in raw (unprocessed) NIFTI formats. Depending on the nature of the EPI functional acquisition, the nine datasets were processed with slightly different preprocessing pipelines. We first describe dataset-specific preprocessing, and then describe the common preprocessing steps that were applied to all datasets following dataset-specific preprocessing. All preprocessing steps were performed in either *Nipype* (Gorgolewski et al., 2011) with calls to *FSL* utilities.

#### Dataset Specific Preprocessing

For the ME-REST, ME-TASK, ME-TASK-CUE and ME-REST-SUPP datasets, multi-echo EPI functional preprocessing pipeline consisted of the following steps: the first 7 (ME-REST, ME-TASK, ME-TASK-CUE) and 4 (ME-REST-SUPP) volumes were removed, six-parameter rigid body motion correction and slice timing correction with *MCFLIRT* and *slicetimer* in FSL, respectively, Multi-Echo ICA denoising and optimal combination of echos using Tedana software (DuPre et al., 2020), non-linear registration to the MNI152 template using FSL’s FMRIB’s Nonlinear Image Registration Tool (FNIRT).

For the HCP-REST dataset, we used EPI functional scans previously preprocessed with the HCP’s ICA-based artifact removal process (Smith et al., 2013) to minimize effects of spatially structured noise in our analysis. EPI scans were previously motion-corrected, registered to the MNI152 template, and intensity normalized. Comprehensive details of the HCP preprocessing pipeline are described in Glasser et al. (2013).

The NKI-REST, NKI-TASK, YALE-REST, NATVIEW-REST and clamped CO2 datasets shared a largely similar set of preprocessing steps. First, the first N volumes (∼10s) of the NKI-REST (7), NKI-TASK (7), YALE-REST (10), and clamped CO2 (26) EPI functional scans were dropped to remove non-steady state time points. For the NATVIEW-REST dataset, the last 2 volumes were dropped to align with the EEG and pupillometry recordings (no volumes were dropped from the start of the scan). Slice timing correction was applied to the EPI functional scans of the NATVIEW-REST dataset using the *FSL slicetimer* utility due to its longer TR acquisition (TR=2.1s). Second, EPI functional scans from all datasets were motion corrected with six-parameter rigid body alignment with FSL *MCFLIRT.* EPI functional scans were then non-linearly registered to the MNI152 template using FSL’s FMRIB’s Nonlinear Image Registration Tool (FNIRT) (Andersson et al., 2010).

#### Common Preprocessing

Following dataset-specific preprocessing, all datasets were then resampled to 3mm (isotropic) MNI152 space. Excluding the HCP-REST dataset, this resampling step occurred during FNIRT registration. For the clamped CO2 dataset, fMRI volumes were resampled to 4mm (isotropic) MNI152 space to more closely match its spatial sampling. Following resampling, fMRI volumes were spatially smoothed with a gaussian kernel (FWHM=5mm) and temporally filtered with a fifth-order Butterworth bandpass zero-phase filter (0.01-0.1Hz). For all datasets, voxels were extracted with a dilated MNI152 brain mask, so as to pick up voxel time courses in large dural venous sinuses and CSF compartments.

#### EEG Preprocessing

Preprocessing of EEG recordings in the ME-REST, ME-TASK and ME-TASK-CUE datasets are described in (Goodale et al., 2021). Briefly, channel time courses were first corrected for gradient artifacts through the average artifact subtraction method (Allen et al., 2000). Ballistocardiogram artifacts were removed by subtraction of an average artifact template locked to cardiac R-peaks, followed by independent component analysis (ICA) of the template-subtracted time courses. The scalp EEG channel time courses were referenced to channel FCz for analysis.These EEG preprocessing steps were performed with the BrainVision Analyzer software.

Preprocessing of EEG recordings for the NATVIEW-REST dataset was performed with custom EEGLAB MATLAB scripts provided by the authors of the dataset (https://github.com/NathanKlineInstitute/NATVIEW_EEGFMRI) and are described in (Telesford et al., 2023). Briefly, the preprocessing steps included 1) gradient artifact removal with the FASTR utility from the EEGLAB FMRIB Plugin, 2) QRS/heartbeat detection using the ECG channel, followed by pulse artifact correction with a template subtraction method, and 3) bandpass filtered between 0.3 to 50Hz using a Hamming windowed sinc FIR filter. Scalp EEG time courses were referenced to the common average for analysis.

#### Physiological and EEG Feature Extraction

Five physiological signals were extracted from the raw PPG, respiration belt, skin conductance, and pupil diameter recordings. Heart rate variability and systolic peak amplitude (PPG amplitude). Respiratory volume (Harrison et al., 2021) was extracted from the raw respiration belt recordings. Low-frequency (0.01-0.1Hz) tonic skin conductance was extracted from the raw skin conductance signals. We used minimally-preprocessed pupil diameter signals extracted from eye-tracking recordings provided by the authors for the NATVIEW-REST and YALE-REST datasets. Example signals from each dataset are displayed in **Supplementary Figure 6**. For comparison with fMRI signals, the extracted physiological signals were clipped at five standard deviations from the mean (for outlier removal), resampled to the length of the functional MRI scan and filtered using a fifth-order Butterworth bandpass filter (0.01-0.1Hz) excluding the tonic skin conductance signals that were already filtered. Note: the minimally-preprocessed pupil diameter signals from the YALE-REST dataset were already resampled to the length of the functional scan. Details of the preprocessing for each physiological signal are provided below. The preprocessing pipeline for the physiological signals (and EEG signals) is illustrated in **Supplementary Figure 8.**

Heart rate variability time courses were extracted from PPG time courses using the NeuroKit2 package (https://neuropsychology.github.io/NeuroKit/index.html) in Python. For calculation of HR, the raw PPG time course was first filtered with a third-order Butterworth bandpass filter (0.5 - 8Hz) followed by systolic peak detection using the method by Elgendi et al. (2013). Heart rate was calculated from the period of time between peaks and interpolated to the same length of the raw signal with monotone cubic interpolation (Fritsch & Butland, 1984). For extraction of PPG pulse amplitude signals, the amplitude of the systolic peaks (previously identified by the peak detection method) were interpolated with monotone cubic interpolation.

For the NKI-REST and NKI-TASK dataset, skin conductance (SC) signals were collected from the hand. SC time courses consist of a low-frequency tonic and high-frequency phasic component (Lykken & Venables, 1971). The tonic component reflects the slowly-varying component of the skin conductance signal, and has previously been studied in the context of fMRI (Nagai et al., 2004). We extracted a narrowband tonic SC signal matching the frequency content of spontaneous resting-state fMRI signals (0.01-0.1Hz) with a fifth-order Butterworth bandpass filter (0.01-0.1Hz).

For the YALE-REST dataset, we used minimally preprocessed pupillometry signals provided by the authors (https://openneuro.org/datasets/ds003673/versions/2.0.1). The minimal preprocessing pipeline consisted of 4-point spline interpolation of eye blinks, low-pass filtering with a Butterworth filter (< 0.5 Hz), removal of the first 10 seconds of recordings (to match the length of the functional scan), and resampling to the sampling frequency of the functional scan (1 Hz). We also used minimally preprocessed pupillometry signals for the NATVIEW-REST dataset. The minimal preprocessing pipeline consisted of linear interpolation of eye blinks, median filter of 0.2 secs width, and resampling to the frequency of the functional scan (0.47Hz).

Respiratory volume (RV) was calculated from respiration belt time courses using a recently developed Hilbert-based method (Harrison et al., 2021), implemented in NeuroKit2.

High-frequency noise was first removed from the respiratory belt time courses with a Butterworth tenth-order low-pass filter (< 0.75Hz). Amplitude and phase components were then extracted from the filtered signal via the Hilbert Transform. Following an iterated linear interpolation procedure of the phase time courses, RV was calculated as the product of the derivative of the interpolated phase time course (i.e. the instantaneous breathing rate) with the signal amplitude (breathing depth/amplitude).

EEG power and vigilance fluctuations were extracted from averaged parietal and occipital lobe EEG channel time courses (ME-REST/TASK: P3, P4, Pz, O1, O2, Oz; NATVIEW-REST: P3,P4, P7, P8, Pz, POz, P1, P2, PO3, PO4, P5, P6, PO7, PO8, O1, O2, Oz) using the MNE-Python package (Gramfort et al., 2013) (https://mne.tools/stable/index.html). Time-frequency EEG power was extracted via Morlet wavelet filters (number of cycles = 15) to construct a filter bank ranging from 2 to 20Hz (spanning Delta, Theta and Alpha oscillation bands). For the ME-REST and NATVIEW-REST dataset, power was extracted from each signal in the filter bank and was cross-correlated with time courses from the first principal component of the fMRI data (see below). Alpha power signals were computed through band-pass FIR filtering (Hamming window; 8 - 12Hz) of the average channel time course, followed by extraction of instantaneous amplitude via the Hilbert Transform. For comparison with fMRI time courses in the ME-REST and NATVIEW-REST dataset, EEG power time courses were resampled to the length of the fMRI scan. Example EEG power time courses from a sample of subjects in the ME-REST and NATVIEW-REST datasets are displayed in **Supplementary Figure 7**.

#### K-Complex Annotation

Event-related averaging of fMRI and physiological signals around K-complex onsets was performed on a subset (N=7) of participants from the ME-REST dataset who fell asleep during their scanning session. K-complex annotations were performed in a semi-automated fashion.

First, EEG time courses from frontal and central channels were automatically sleep staged using the yasa package in Python (https://github.com/raphaelvallat/yasa; Vallat & Walker, 2021) with default parameters. Following automated sleep-staging, EEG time courses from sleep stage II were manually annotated for K-complex onsets according to AASM criteria (Berry et al., 2017) using a custom bipolar montage. Separate K-complexes that occurred within a window of ∼30s were removed before event-related averaging to avoid overlapping responses.

#### Principal Component Analysis

As in Bolt et al. (2022), the zero-lag and time-lag structure of the global fMRI signal was modeled with principal component analysis (PCA) and complex principal component analysis (CPCA), respectively. As shown in Bolt et al. (2022), the first principal component of both PCA and CPCA extract a pattern of global fMRI fluctuations that is closely correlated (in time) with the global mean time course. Standard PCA was used to extract the simultaneous statistical dependence between cortical areas of global fMRI fluctuations, CPCA was used to extract time-lag statistical dependence (i.e. traveling wave or propagatory behavior). In this study, we analyze the properties of the first principal component in volume space, including signals from subcortical structures, ventricles and cerebral sinuses.

CPCA involves the application of PCA to complex-valued time courses generated by the Hilbert transform. We extracted the first complex principal component using CPCA on the complex-valued (bandpassed to 0.01-0.1Hz) fMRI time courses temporally-concatenated across subjects. Time-lag information can be extracted from the phase representation of the complex-valued PC (via Euler’s Identity). The phase representation encodes the time-delay between voxels (in radians) within the first complex PC. The time-lag information can also be visualized over selected time points via a temporal reconstruction (**Supplementary Movie 1**). Comprehensive details are provided in Bolt et al. (2022). Briefly, we first divided the temporal phase time courses of the first complex PC into equally-spaced bins (N=30). We then projected the complex PC back into voxel space to derive voxel time courses. Finally, we averaged the real-valued voxel time courses within the time points indexed by the equally-spaced phase bins. This resulted in a 30-volume ‘movie’ that visualizes the temporal evolution of a component. Both PCA and CPCA solutions were computed using a fast randomized SVD algorithm developed by *Facebook* (https://github.com/facebookarchive/fbpca). More details of the CPCA algorithm can be found in Bolt et al. (2022).

#### Cross-Correlation Analyses

For the ME-REST, HCP-REST, ME-REST-SUPP, NKI-REST and YALE-REST datasets, cross-correlation analyses were conducted between the electrophysiological time courses and the first principal component (PC1) time course. Product-moment cross-correlations were computed at the subject-level and group-average cross-correlations were computed by the mean of the subject-level cross-correlations. For comparability across datasets with different sampling rates, we interpolated subject and group-average cross-correlation functions with a cubic spline from -30s to 30s (time-lag).

#### Impulse Response Modeling of Physiological Signals

To summarize the temporal dynamics of all recorded physiological signals in response to fluctuations in global fMRI signals (i.e. PC1), we implemented a linear systems modeling approach as described in Chang et al. (2009). Impulse response functions were estimated separately for each physiological signal through deconvolution of the physiological signal using a Gaussian Process prior. A Gaussian Process prior is used to capture the underlying smoothness of the physiological signal. The deconvolved physiological signal is estimated from the maximum *a posteriori* (MAP) of the posterior distribution. As no single dataset contained all physiological signals, we superimposed the deconvolved physiological impulse responses from three separate datasets: ME-REST (PPG signals, Alpha EEG power, and respiratory volume), NATVIEW-REST (pupil), and NKI-REST datasets (skin conductance). The length scale and kernel variance hyperparameters of the Gaussian Process, were set to 3 and 1, respectively.

Negative shifts of the signal were included to ensure that physiological responses before the peak of the global fMRI signal were captured.

#### Multi-Set Canonical Correlation Analysis

To estimate the *joint* fluctuations between physiological and the first principal component time courses, we implemented a multi-set canonical correlation analysis (MCCA) on the full set of signals in each dataset (Kettenring, 1971). Each ‘set’ was formed from time lags of a single physiological signal. Inclusion of all potential time lags out to a window size N leads to potential collinearity and risks overfitting. Instead, we implement a generalized additive distributed lag approach (Gasparrini et al., 2010; Zanobetti et al., 2000), where time-lagged predictors for each physiological signal were generated via a linear combination of time-lagged copies of the signal with a natural cubic spline basis with three splines distributed across the time window (from 0 to 10s; number of time points varied for each dataset due to different sampling rates). The resulting time-lagged predictor is represented by three linear-weighted versions of the original signal encoding a smooth curve across the time window.

The objective of MCCA is to find a linear-weighted combination of the time-lagged copies of each signal that maximizes the pairwise correlations between all signals. MCCA was performed at the group-level by group-wise temporal concatenation. The number of ‘sets’ in the MCCA analysis varied across datasets due to the differing number of physiological signals recorded across datasets. We extract the first canonical component from the MCCA algorithm, corresponding to the linear weighted combination of all signals that produces the maximum pairwise correlation between signals. The MCCA algorithm was implemented in the cca-zoo Python package (https://github.com/jameschapman19/cca_zoo). Illustration of the MCCA approach used in our study is provided in **Supplementary Figure 9**.

Statistical significance testing of the average pairwise correlation of the first canonical component was performed using block-wise permutation of subject time courses before temporal concatenation across subjects. Specifically, physiological signals were randomly shuffled across subjects before temporal concatenation, effectively removing cross-signal couplings while preserving the autocorrelation structure within each signal. For example, one permutation may align signal A in subject 1, with signal B in subject 7, and so forth. For each permutation (N=1000), the average pairwise correlation between all permuted time courses of the first canonical component was extracted to construct a null distribution.

#### Event-Related Averages

Event-related averaging of the ME-TASK, NKI-TASK and ME-REST datasets was performed to examine the fMRI, EEG and physiological response to deep breaths, breath holds and spontaneous K-complex onsets, respectively. A group-level response function for each physiological signal was generated by averaging across all trials and subjects within a window starting from onset out to ∼29s. Standard error plots for each group-level response function were generated via cluster bootstrapping, where subjects were randomly resampled with replacement (N = 100) before averaging.

For the ME-TASK dataset, event-related EEG power fluctuations were examined for each subject by averaging Wavelet filter bank power signals (i.e. time-frequency spectral power in 2-20Hz frequencies; see above) within a 20s window post breath-onset across trials. Baseline log-ratio normalization (i.e. decibels) was applied to the subject-averaged time courses with the time span 1s before up until stimulus onset as the baseline. A group event-related average was constructed from averaging across event-related subject averages.

For the ME-TASK-CUE dataset, event-related averaging of physiological and EEG power time courses was conducted using the same procedure as the ME-TASK dataset. Some inter-trial intervals between auditory cues were shorter for this task, on average (range: ∼8 to 89 secs), relative to to the ME-TASK dataset. Trials separated by less than 30s were excluded. In addition, trials with a non-response (i.e. no button response) were excluded.

#### Multi-Echo Modeling of T2* and S0 Effects

To explore potential hemodynamic effects in the global fMRI signal we modeled T2* decay rates and initial signal intensity (S0) signals from the ME-REST-SUPP dataset. The T2* and S0 time signals were estimated from three consecutively acquired echos at each time point via a log-linear least squares fit to a monoexponential curve, 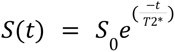. The volume-wise log-linear least squares fit of the consecutively acquired echos to a monoexponential curve was performed with the *tedana* Python package (DuPre et al., 2020) with the *fit_decay_ts* module.

#### Cardiac RETROICOR Regression of Multi-Echo ICA Denoised Signals

Variations in the cardiac cycle are known to induce pulsatile flow changes in cerebral arteries causing fMRI signal fluctuations and other artifacts (Dagli et al., 1999). Our analyses showed that variations in heart rate were associated with global fMRI fluctuations (**Figure 1B**), suggesting that cardiac-driven pulsatile flow changes may be prominent during global fMRI fluctuations. However, we expected artifactual fMRI signal variations from vessel pulsation and tissue deformation due to the cardiac cycle to be largely attenuated in ME-ICA denoised signals.

To demonstrate that pulsatile blood flow fluctuations of the cardiac cycle do not significantly contribute to ME-ICA denoised signals, we performed regression modeling with a Fourier series expansion of cardiac cycles acquired from PPG signals (RETROICOR regressors; Glover et al., 2000). Because slice-timing correction was performed before ME-ICA denoising, RETROICOR cardiac regressors estimated at each slice (to account for potential differences in slice timing) were regressed separately onto all voxel denoised signals, and the maximum explained variance (R2) across slice-specific regressors was selected (**Supplementary Figure 12**). To demonstrate that pulsatile-driven fMRI signal fluctuations were isolated and rejected as ‘noise’ components by the ME-ICA algorithm, we also regressed

RETROICOR cardiac regressors onto ‘noise’ signals constructed from rejected ICA components.

